# Transcriptional noise sets fundamental limits to decoding circadian clock phase from single-cell RNA snapshots

**DOI:** 10.1101/2024.06.30.601408

**Authors:** Anjoom Nikhat, Taniya Mandal, Nivedha Veerasubramanian, Shaon Chakrabarti

## Abstract

The rhythmic circadian clock generates temporal variation of critical physiological processes in tissues, making accurate measurement of clock phase or ‘tissue time’ a fundamentally important problem. Recent advances in single-sample time inference provides a potentially powerful alternative to traditional time-series based approaches. Single-sample techniques typically leverage population-level RNA rhythms, but the feasibility of single-cell phase detection remains an open question. Combining multiplexed smFISH to simultaneously measure up to 6 mouse fibroblast genes, with a novel inference algorithm using Gaussian Processes, here we demonstrate that even when technical drop-outs are minimized, transcriptional noise in core-clock genes precludes single-cell phase inference. Simulations predict that above 50 ‘clock-like’ genes would make single-cell phase inference possible. Remarkably however, just 3 core-clock genes are sufficient if RNA levels of ∼ 70 cells are averaged. Further, we demonstrate how averaging allows detecting spatially-resolved, heterogeneous clock phases in desynchronized cells. Our work provides a conceptual framework for achieving high-resolution phase detection with a minimal set of core-clock genes, with implications for probing the origins of clock dysfunction, otherwise unresolvable using population-based approaches.

## Introduction

Internal clocks present across organisms have evolved in the presence of a rhythmic external environment to allow anticipation of external time. At least one of these time keepers, the circadian clock, is endogenous and operative at the level of single cells. At the molecular level, a set of transcriptional and translational feedback loops (TTFLs) combine to generate approximately 24 hour oscillations of a set of “core” genes and proteins [1]. These molecular clocks in the Suprachiasmatic Nucleus (SCN) of the mammalian brain get entrained to the external light-dark cycles, resulting in an accurate internal timer with a 24-hour time period. The SCN in turn largely controls entrainment of the millions of cells constituting various peripheral tissues, thereby providing a notion of internal time across the body [2]. Inferring the internal time (alternatively referred to as ‘circadian phase’) is of immense importance in chronobiology, since many critical physiological processes are timed by the circadian clock [3, 4, 5], and hence could be utilized for chronotherapy [6, 7, 8, 9, 10, 11, 12]. Most methods rely on measuring a proxy of the molecular clock, the “chronotype”, either by using qualitative questionnaires on an individual’s sleep and meal patterns [13], or more quantitatively by biochemically detecting the onset of melatonin [14]. Recent approaches have used regression techniques based on time series data generated by wearable devices, to predict melatonin onset and internal clock phase [15]. However, issues of accuracy, logistical challenges and masking by environmental factors have spurred recent interest in developing alternate and easier approaches to directly measuring the molecular clock phase. With rapid advances in high throughput gene expression assays and concomitant development in Machine Learning algorithms, single sample inference of circadian phase has emerged as an exciting area of research [16]. State-of-the-art methods typically use either supervised or unsupervised learning to predict the circadian phase of easily accessible tissues (like blood or skin), using RNA measurements of ∼ 10 − 15 core-clock genes averaged over millions of cells [17, 18, 19, 20, 21, 22, 23, 24, 25, 26, 27, 28].

Since single sample phase inference techniques till date have almost solely focused on utilizing bulk RNA technologies, very little is currently understood about the possibility of phase inference from *single-cell* RNA snapshots. If possible, this would open up a new and exciting frontier of circadian phase inference – allowing for discovery of spatial heterogeneity in endogenous clock phase within single tissue samples, determination of the underlying cause of circadian dysfunction and dampened rhythms in diseases like cancer, and also allowing for the possibility of investigating mechanisms of inter-cellular coupling leading to phase (de)coherence in peripheral tissues. Historically, single-cell circadian clock measurements have been performed using promoter-driven luciferase reporters in mouse tissue explants [29, 30], or knock-in reporters of proteins [29, 31, 32]. These methods however are limited to model systems due to the necessity of generating genetically modified organisms, and/or do not quantify the endogenous levels of the clock RNAs. While scRNA-seq has recently been used to develop phase inference methods like Tempo [33] and tauFisher [34], associated technical difficulties necessitate “pseudo-bulk” analyses for accurate circadian phase inference, typically requiring averaging over hundreds of cells [33, 34, 35]. Although Tempo and tauFisher improve over older algorithms like Cyclops [24] and Cyclum [36], they still give large errors in phase inference at the single-cell level. Whether this arises due to intrinsic gene-expression variability of the clock genes ([37]) or from technical noise invariably associated with scRNA-seq measurements, remains an open question. Crucially, since scRNA-seq loses direct spatial information, it limits our ability to study phase variation across space. Some studies have utilised single-molecule FISH to obtain more accurate measurements of endogenous RNA in mouse SCN and liver, but they have been limited to a couple of clock genes and do not address the question of possibility of circadian phase inference from single cells [38, 39].

In this study we systematically probe the limits of single-cell circadian phase decoding from RNA measurements of core-clock genes. To circumvent the major technical limitations of single-cell sequencing approaches, we utilise a recently developed multiplexed smFISH technique SABER-FISH [40] to obtain absolute RNA counts of up to 6 core-clock genes simultaneously in hundreds of mouse embryonic and lung fibroblasts. To optimally analyze the single-cell data, we use Gaussian Processes to develop a supervised phase inference algorithm PRECISE (**PRE**dicting **CI**rcadian phase from **S**tochastic gene **E**xpression). Using PRECISE as well as a variety of other unsupervised algorithms, we demonstrate that biological, not technical noise, in these measured genes prevents inference of circadian phase from single cells. Systematically adding more genes in computer simulations, using noise levels similar to the least noisy core-clock gene *Nr1d2*, allows for accurate single-cell phase inference only above 50 genes. Given that there are only about 10-20 genes of the circadian clock network that are consistently found to oscillate across tissues, our results suggest that it may not be possible to decode single-cell clock phase just from core-clock genes. Remarkably however, averaging over as few as ∼ 70 cells allows for accurate phase inference with ∼1 hour error using just 3 core-clock genes. Finally we demonstrate how a strategy of coarse-graining over a small group of spatially proximal cells can be used to resolve heterogeneous circadian phases within a single asynchronized population. Our results suggest that while single-cell circadian phase inference from core-clock RNA snapshots may be fundamentally limited by transcriptional noise, coarse-graining over small groups of cells may still allow for spatially resolved phase inference at near single-cell resolution.

## Results

### A quantitative definition of circadian clock phase from population level RNA measurements

The oscillatory nature of core-clock gene transcripts of a synchronized population of cells, combined with the well characterized network of genes forming the circadian clock, allow for a quantitative definition of circadian clock phase. Cells at distinct times post synchronization represent distinct circadian phases (Figure 1A left), with previously well characterized functional differences. The time post synchronization can be uniquely identified by the relative (average) RNA levels of a group of core-clock genes, which oscillate with fixed phase relationships between each other (Figure 1A left) [1]. These times can equivalently be described by their angular position along the high-dimensional ellipse formed in gene expression space, which justifies the equivalent usage of the term ‘phase’ for the times post synchronization (Figure 1A right). Note that any of the oscillatory core-clock genes can be used to define the zero phase angle. If not mentioned otherwise, throughout this work we use the peak of the *Nr1d1* gene to define circadian phase 0 for NIH3T3 cells and the peak of the *Tef* gene to define circadian phase 0 for MLG cells.

**Figure 1:**
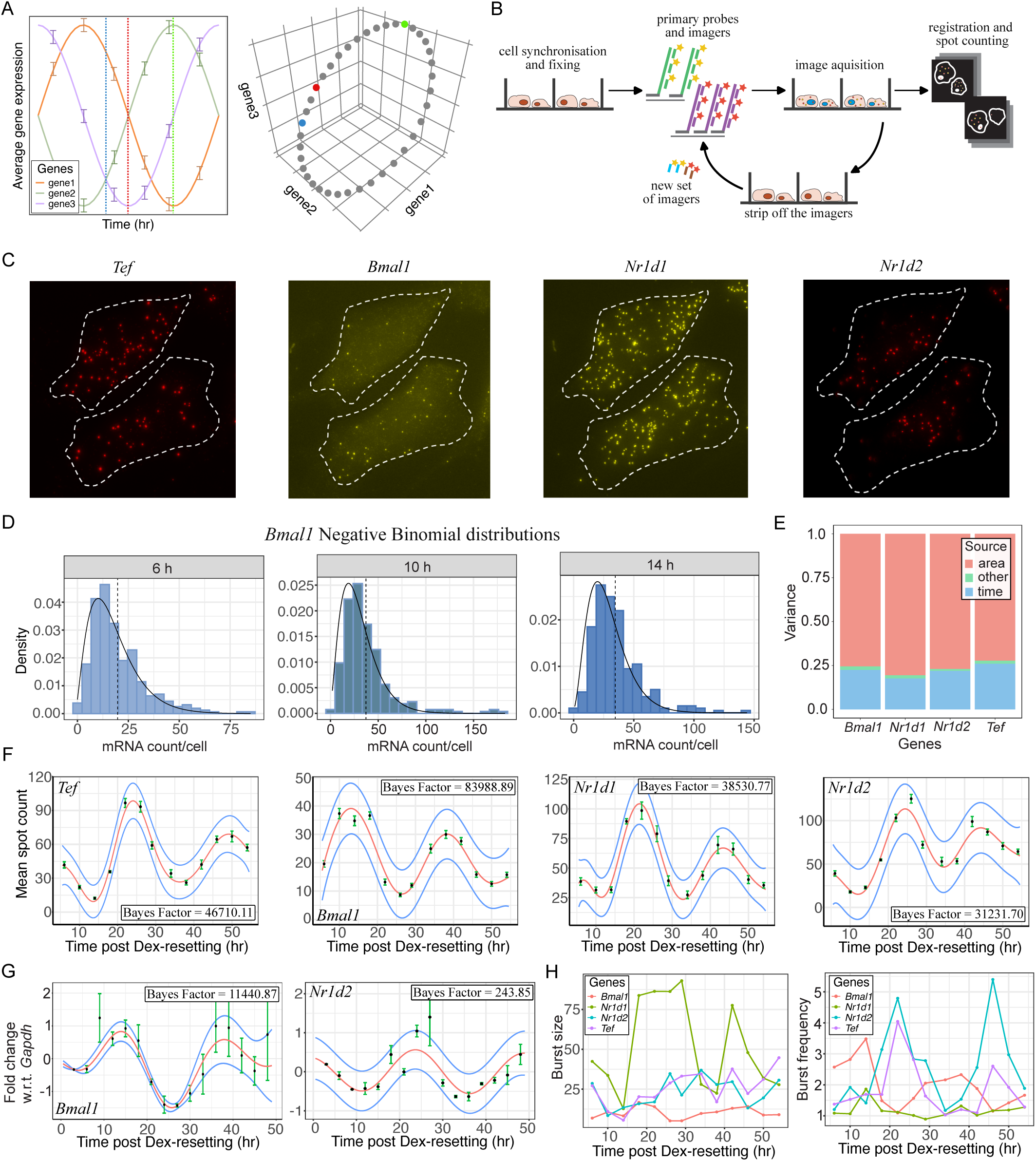
Single-cell gene expression recapitulates bulk properties and defines phases of the circadian clock. (A) Schematic showing how the relative levels of average gene expression uniquely defines circadian phases.. The blue, red and green vertical lines correspond to 3 distinct circadian phases (left) and are positioned at unique angles on the high dimensional ellipse (right). (B) Schematic of the SABER-FISH imaging process. (C) An example of SABER-FISH RNA spots for the 4 clock genes, shown in the same two cells. (D) Negative Binomial fits to *Bmal1* RNA counts at three times post Dex resetting, showing a clear shift in the mean. (E) Variance decomposition with area and time as covariates shows a significant contribution from time. (F) Mean RNA counts and errors in the mean as functions of time. Red lines – ODeGP [43] posterior mean, blue lines – posterior variance. Bayes Factors larger than 14 imply statistically significant oscillations. (G) Bulk qPCR on millions of cells showing the same oscillatory patterns and phase differences for *Bmal1* and *Nr1d2* as observed in panel F. (H) Both burst size and burst frequency show oscillations over time.

A crucial aspect of this definition of circadian phase based on the core-clock RNA levels is the averaging over synchronized cellular populations. Previous algorithms have used this definition to infer circadian phase from bulk gene expression datasets [17, 18, 19, 20, 21, 22, 23, 24, 25, 26, 27, 28]. What remains unanswered is whether the same relative RNA levels, as defined by the population averages, can be used to ascribe correct circadian phases to single cells. Answering this question is the central goal of this work.

### Single-cell endogenous RNA imaging recapitulates bulk oscillations and defines phases of the circadian clock

To generate cells in these different circadian phases, we used Dexamethasone (Dex) resetting to synchronize NIH3T3 cells and then captured the cells at a few discrete times post-Dex for multiplexed RNA measurements. To reduce technical noise in the measurements as far as possible, we used the recently developed SABER-FISH (Signal Amplification By Exchange Reaction - Fluorescence in situ Hybridisation) method to measure single-cell endogenous RNA counts [40]. This method uses long repetitive sequences called ‘concatemers’ at the end of probes complementary to the RNA sequences of interest, which then act as platforms to hybridise many fluorescently labeled ‘imagers’ thus amplifying the signal from each RNA molecule [40, 41]. Besides the signal amplification allowing for sensitive detection of low expression genes, SABER-FISH additionally allows for multiplexing to simultaneously measure multiple genes in the same single-cell (Figure 1B). To validate the method, we first imaged a well characterized gene *Cbx5* [40] in the two cell lines used in this study – NIH3T3 and MLG. We consistently achieved ∼ 95% co-localization in both cell types upon using two imagers simultaneously to image *Cbx5* RNA spots (Figure S1, details in Section S1), thereby demonstrating the accuracy of the measurements.

To collect cells from distinct circadian phases, we followed a protocol of adding Dex at different times and fixing cells simultaneously to minimize cell density variation amongst different time points (Figure S2, details in Section S1). After fixation, we performed multiplexed SABER-FISH to obtain absolute RNA counts of the 4 core-clock genes *Bmal1*, *Tef*, *Nr1d1* and *Nr1d2*, in about 200-250 cells per time point (Figure 1C; experimental details in Methods and Sections S1 and S2). We first checked that the distributions of RNA counts for each gene across single cells followed the Negative Binomial distribution (details in Section S3) – a well established signature of transcriptional bursting (Figure 1D shows the distributions for *Bmal1*; other genes are shown in Figure S3). Given the large ‘super-Poissonian’ noise describing the expression distributions *within* each time-point, we next asked whether expression variation *across* time-points could be detected. We therefore performed variance decomposition analysis [42] (details in Section S3) and found that a significant 20-25% of the total variance could be ascribed to time (Figure 1E), suggesting that average levels of gene expression across time are likely to be well separated despite the high intrinsic noise. To confirm this expectation, we bootstrapped the data from each time point to generate mean expression levels and errors of the mean (Figure 1F). The mean expression for all 4 clock genes showed strong oscillations when analyzed with ODeGP (Figure 1F; details in Section S3), an oscillation detection algorithm that we previously developed and demonstrated to outperform existing methods [43]. Indeed, the oscillatory patterns recapitulated well known phase differences between these four core-clock genes, which we also confirmed for a few genes with qPCR (Figure 1G; details in Section S4). Finally, to identify which parameters underlying transcriptional bursting give rise to the expression oscillations, we extracted the burst size and frequency from the RNA count distributions per time point (see Section S3 for details). We found that both parameters tend to be oscillatory over time (Figure 1H). Therefore, our data suggests that oscillations in both the burst size as well as the burst frequency combine to generate the periodic patterns of gene expression rather than one being constant over time as suggested in previous work [37].

In conclusion, the results of this section confirm that SABER-FISH can be used to accurately measure intrinsic RNA copy number noise among single cells in 4 core circadian clock genes. Recapitulation of strong oscillations in all four genes along with the expected phase differences upon averaging over ∼ 100 cells further demonstrated the accuracy of our datasets, measurement sensitivity and Dex synchronization protocol. Most importantly, these results suggest that the average gene expression levels at different time points post Dex resetting can be used to define unique phases of the circadian clock in NIH3T3 cells.

### Emergence of gene-expression oscillations with expected phase differences upon averaging over **∼** 10 cells

Given that averages over ∼ 100 cells recapitulated known oscillatory patterns and phase differences between the 4 core-clock genes, we next explored whether these oscillations and phase relationships could be observed at the single-cell level. As observed in Figure 1D, the absolute RNA counts of the clock genes showed high variance even within a single time-point post Dex synchronization, suggesting that at the single-cell level it might be challenging to detect oscillations. We created ‘pseudo-time’ trajectories by randomly selecting one cell from each time point and plotting the absolute RNA counts as a function of time (Figure 2A left). We repeated this procedure 50 times to generate 50 such trajectories, depicted by the different colours in Figure 2A (left). If the intrinsic gene expression noise had been low, this procedure should have given rise to similar oscillatory trajectories for all 50 replicates. Instead, the pseudo-time trajectories at the single-cell level were characterised by high noise (Figure 2A left and Figure S4) suggesting that placing single cells along an oscillatory trajectory might prove challenging. Remarkably however, when gene expression was averaged over groups as small as 10 cells per time point, clean oscillations emerged (Figure 2A centre and right) as well as the correct phase differences between the genes (Figure S4), indicating that assigning unique phases along a circadian oscillation might be meaningful only after averaging out the transcriptional noise.

**Figure 2:**
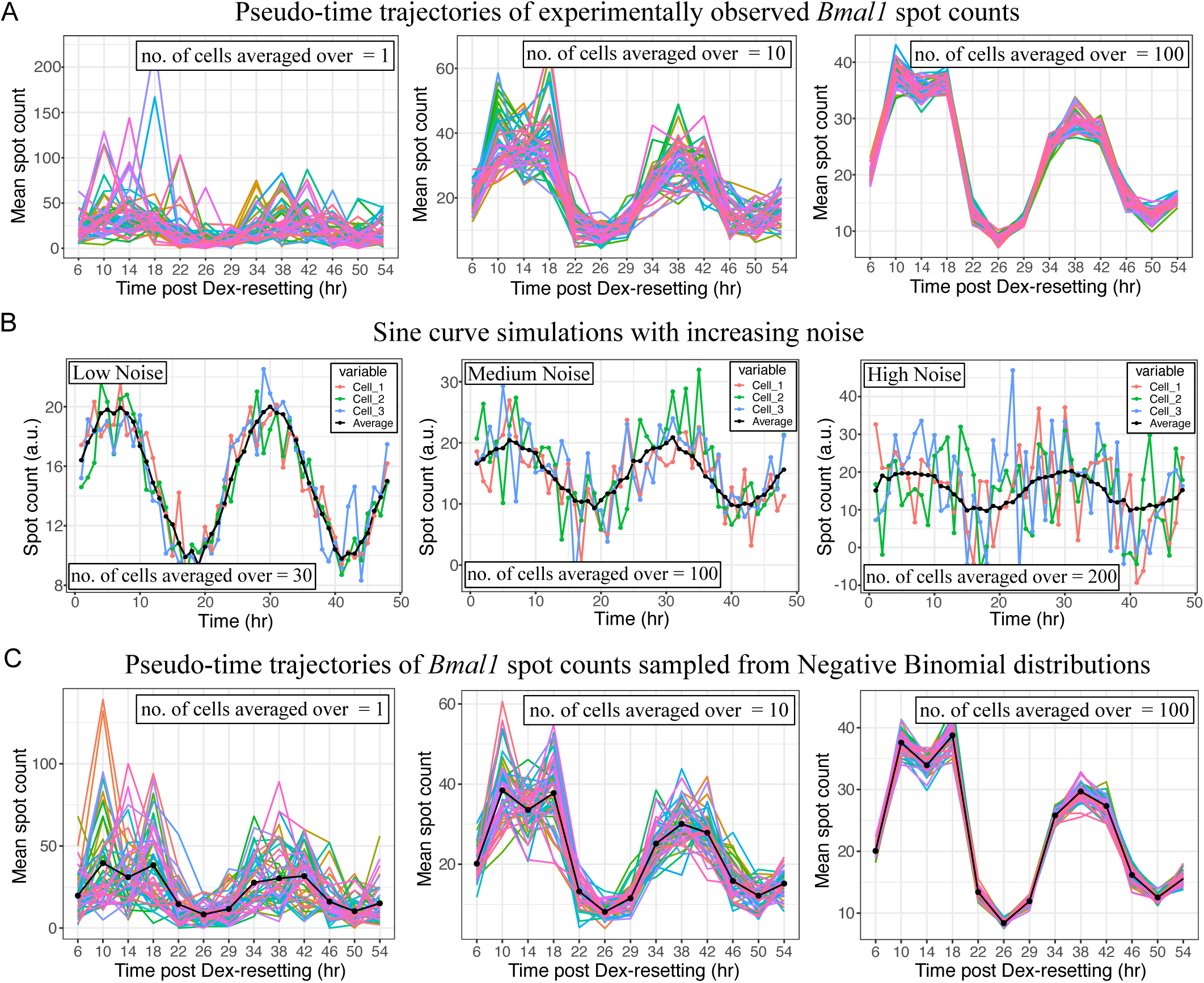
Emergence of oscillations and correct phase differences upon averaging over a small group of single cells. (A) 50 pseudo-time *Bmal1* trajectories of single cells from the experimental dataset (different colours), characterised by high noise (left). Pseudo-time trajectories generated using gene expression values averaged over 10 (centre) or 100 (right) cells demonstrate emergent and low noise oscillations. (B) Simulated sine curves with low (Std. Dev. = 1.5) (left), medium (Std. Dev. = 5)(centre) or high (Std. Dev. = 10) (right) Gaussian noise. Detecting oscillations for the single cells (marked by different colours) is difficult especially in case of high noise. However, oscillatory trajectories emerge upon averaging in all cases (black curves). (C) Pseudo-time trajectories for *Bmal1* generated by sampling from Negative Binomial distributions using parameters obtained from fits to our SABER-FISH datasets from Figure 1D. While single-cell trajectories (left) are characterised by high noise, clear oscillations emerge upon averaging over small cellular groups (centre & right), consistent with our experimental observations.

To gain further intuition on how averaging over cells might lead to reduction in noise and the emergence of clean oscillations as seen in Figure 2A, we carried out simulations using two approaches (details in Section S5): (1) generating time trajectories from a noisy sine function and averaging over many such trajectories and (2) generating time trajectories by sampling from Negative Binomial distributions with burst size and frequency parameters obtained from Figure 1H, and then averaging over many such trajectories. While approach (1) is purely for demonstrative purposes, approach (2) uses parameter estimates from our smFISH datasets, hence is a good representation of the underlying gene expression noise amongst the clock genes. Both methods demonstrate that even if oscillations are hard to detect in single trajectories (coloured lines in Figure 2B-C), averaging removes this noise and produces clear oscillations (black lines in Figure 2B-C). The more the underlying noise level, the more the number of cells required to observe the emergent oscillations, and hence it is interesting that only ∼ 10 cells are sufficient to generate clean oscillations in the experimental dataset (Figure 2A).

In summary, the results of this section suggest that the clock gene expression might to too noisy to robustly assign circadian cell-state or phase to single cells. Intriguingly however, averaging over as few as 10 cells from each time point allowed observation of clear oscillatory patterns along with the expected phase difference between genes, suggesting that averaging over very small groups of cells may be sufficient to overcome intrinsic expression noise and allow accurate assignment of circadian phase.

### PRECISE – an accurate supervised learning algorithm for circadian phase inference

Having developed an intuition for the noise levels in clock gene expression and an approximate estimate of the number of cells required to observe emergent oscillations, we then developed a quantitative approach to establish the limits of circadian phase inference from a single RNA snapshot, in terms of minimum cell and gene numbers required. Since supervised machine-learning approaches tend to outperform unsupervised ones, we first developed a supervised algorithm called PRECISE (**PRE**dicting **CI**rcadian phase from **S**tochastic gene **E**xpression), to accurately infer the circadian phase from RNA counts of a test sample. PRECISE builds on an algorithm we previously developed using Gaussian Processes (GPs) to learn noisy and non-stationary oscillatory trajectories [43]. Here we provide a brief overview of the method; for more details see Methods and Section S6.

The true circadian phases of the training samples are first estimated from the Dex-synchronized time-series of normalized average spot counts (obtained by bootstrapping and normalization of the data), using a wavelet-based approach [44] (Figure 3A and Figure S5). This is followed by learning of the optimum Bayesian posterior mean and variance of the expression levels as functions of phase, for each of the four genes individually. Using the posterior mean and variance, PRECISE then constructs the likelihood of a test dataset characterised by a set of clock gene expression values, as a function of circadian phase (Figure 3A). Finally, PRECISE outputs an average phase estimate (*ϕ_pred_*) determined by detecting positions of local maxima in the likelihood versus phase curve and obtaining their circular mean.

**Figure 3:**
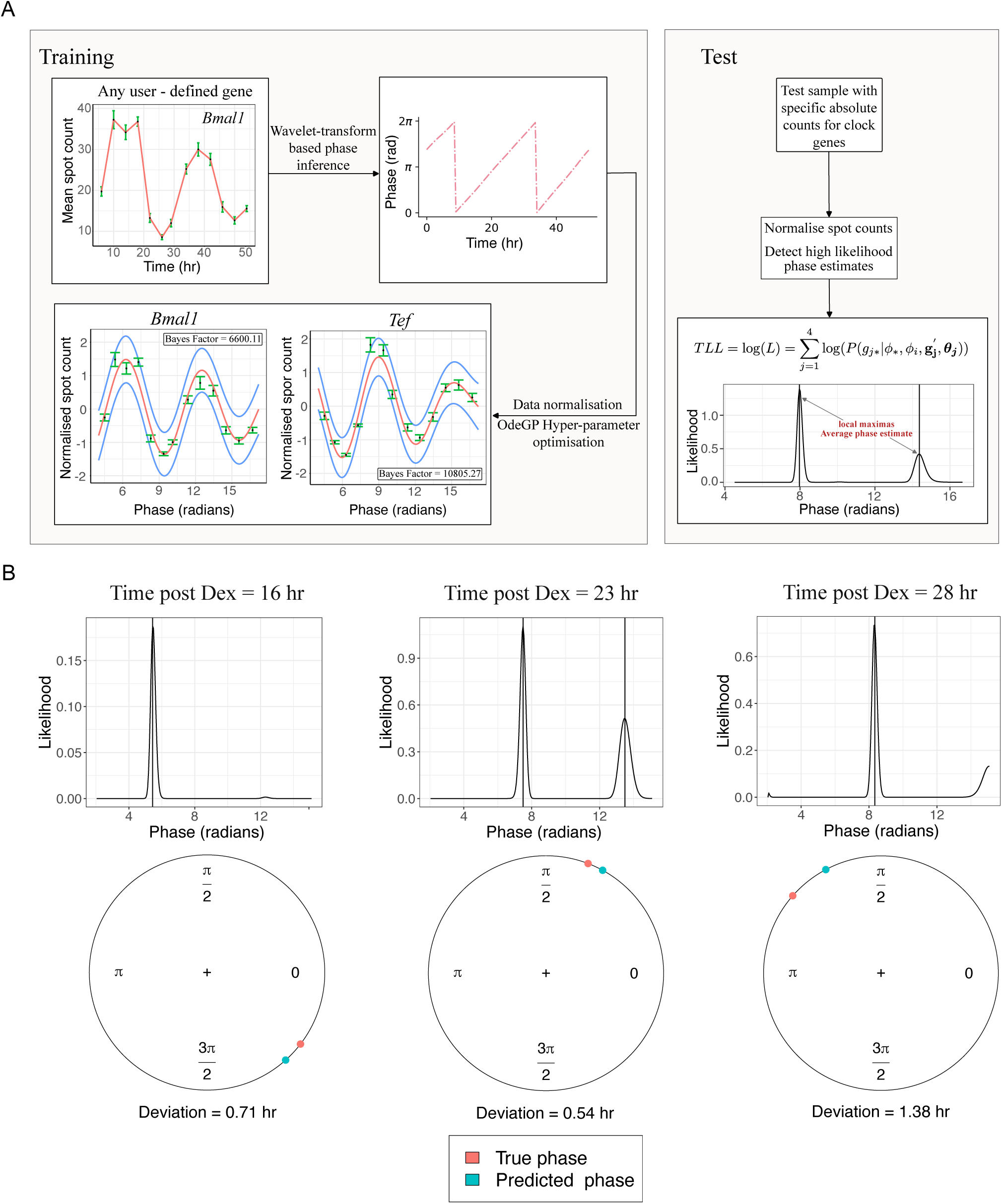
Accurate circadian phase inference using PRECISE. (A) Flowchart indicating the workflow of PRECISE. Training: Mean gene expression at the various times post Dex resetting is first used to assign the circadian phase to the training samples. This is performed using wavelet-transforms. The mean gene expression is then expressed as a function of the circadian phase and normalised. PRECISE then uses Gaussian Process regression to learn the optimised Bayesian posterior mean and variance of the expression levels of the different genes individually. Test: When the raw spot counts of circadian genes for a test sample are provided to pre-trained PRECISE, it normalises the data and constructs the likelihood as a function of the training phases. PRECISE then outputs the predicted circadian phase *ϕ_pred_* of the test sample by calculating the circular mean of the phases corresponding to the likelihood peaks. (B) Validating PRECISE for 3 test time points using gene expression data averaged over ∼ 200-250 cells and 4 genes. Top: Likelihood versus phase curves show clear peaks, allowing accurate phase estimation. Bottom: quantification of deviations between true and predicted phase for the 3 test cases.

For validating PRECISE, we collected SABER-FISH spot count data in Dex-synchronised NIH3T3 cells for 5 test time points not included in the training data (8, 16, 23, 28 and 32 hours). For each time point we imaged ∼200-250 cells, and using pyBOAT assigned the true circadian phase to the 3 centre time points 16, 23 and 28 hours (see Methods and Section S6 for why accurate true phase can be assigned only to the centre time points). Using all 4 genes and averages over the entire cell population, we inferred the test sample phase likelihoods using PRECISE (Figure 3B top). We then computed the predicted phases of the test samples from the likelihood peaks (see Methods for details) and compared the predictions against the true phases (Figure 3B bottom). The deviation Δ between true and predicted phases were on average less than 1 hour (Figure 3B bottom), thus verifying that PRECISE can be used to accurately infer circadian phase with only a few genes.

### Averaging core-clock RNA levels over 20-70 cells is essential for accurate circadian phase inference using PRECISE and SVD

Having validated the ability of PRECISE to accurately infer circadian phase, we then systematically investigated the effect of decreasing the number of genes and cells on the accuracy of phase inference. We used the same test time points – 16, 23 and 28 hours post Dex-resetting and calculated the deviation of PRECISE-inferred phase from the true phase of the cells (Δ; see Methods and Section S6). We first found that using just 2 genes for phase inference usually produced large values of Δ, though in a few cases combinations of two anti-phasic genes worked well (Figure S6). With 3 genes however, accurate phase inference was always possible, though only after averaging over increasing number of cells. As shown in Figure 4A (23 h test sample is shown; other time points are included in the Figure S7), using only the genes *Bmal1, Nr1d1* and *Tef*, the values of Δ almost uniformly covered the full range of 12 hours when single-cell gene expression was used. This suggests that it may not be possible to decode circadian phase with any confidence from single cells. However, upon averaging gene expression over as small a group as 20 cells, PRECISE was able to predict the circadian phase with a median Δ ∼ 1 h (Figure 4A), which is comparable to errors generated by supervised algorithms like ZeitZeiger using gene expression data averaged over millions of cells and at least 10-13 genes [17]. Δ went down further to ∼ 0.5 hours upon averaging over 130 cells, and the variation in Δ shrunk to less than an hour, suggesting that PRECISE is an accurate phase decoder (Figure 4A). We also visualized these deviations of inferred phase using empirical cumulative distribution function (eCDFs), to better understand the performance of PRECISE. The eCDF using single cells was close to the diagonal line (red dotted line in Figure 4B), suggesting that predictions using single cells are only slightly better than making purely random choices of phase. For averages over ≥ 20 cells, the eCDFs were close to optimum with minimum performance improvement for higher sample sizes, further emphasizing that gene expression when averaged over small cellular groups allows for accurate phase estimation (Figure 4B). Finally we compared Δ when different combinations of 3 genes were used as well as when all 4 genes were used (Figure 4C). While all gene sets yielded Δ close to 1 h, interestingly 3 clock genes allowed phase inference as accurately as 4 genes (Figure 4C), suggesting that the phase information provided by gene expression likely saturates at only 3 genes. All these results were consistent irrespective of which gene was used to assign true phase and the time point analyzed (Figure S8).

**Figure 4:**
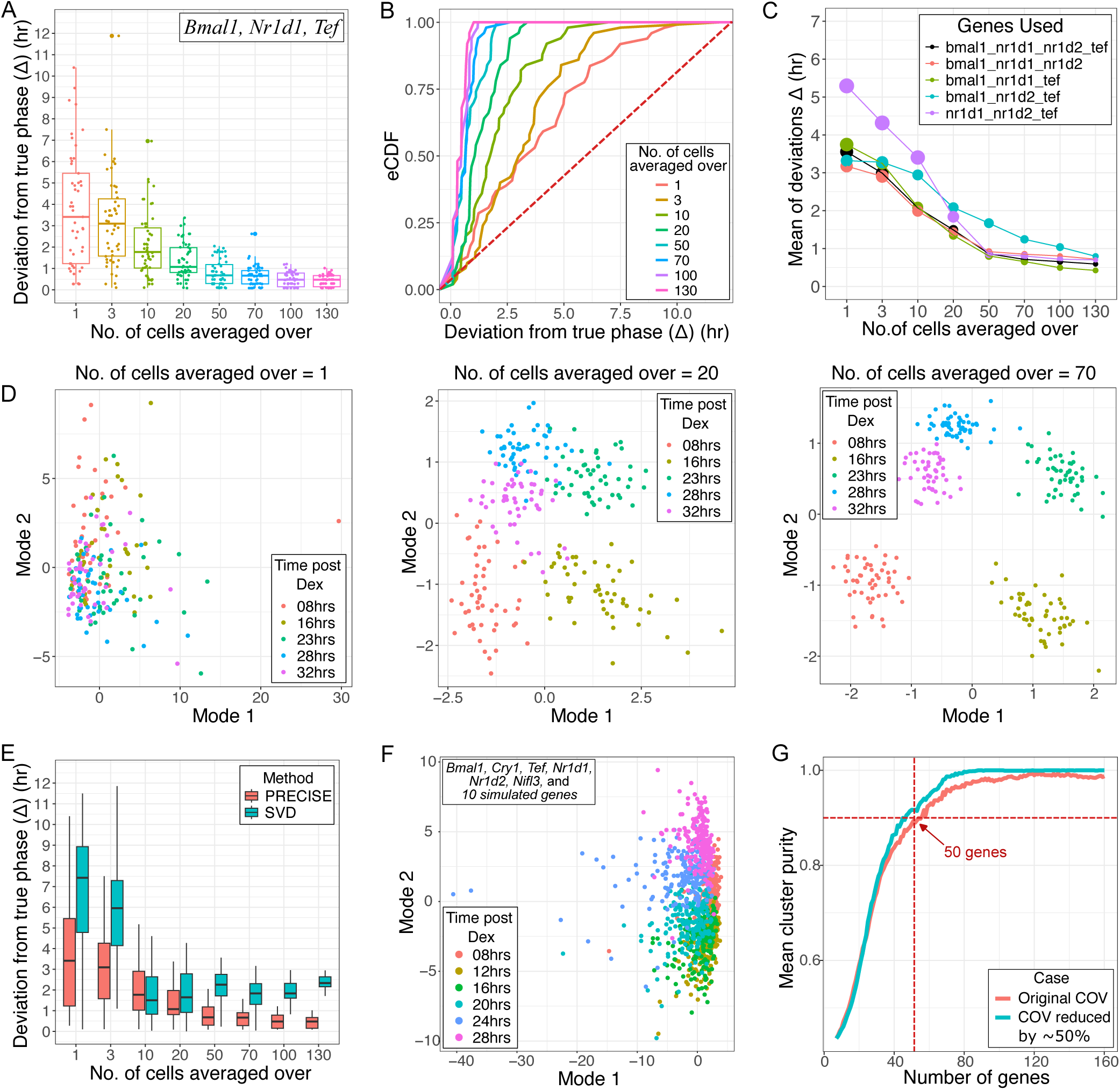
Coarse-graining over small cellular groups is essential for accurate phase inference from core-clock RNA levels. (A) Deviation of PRECISE inferred phase (*ϕ_pred_*) from the true phase assigned when genes *Bmal1, Nr1d1* and *Tef* are used. Accurate phase inference is possible when expression values are averaged for small cellular groups *>* 20 cells as characterised by median Δ ∼ 1 hr. (B) Empirical cumulative distribution functions (eCDFs) for Δ. The single-cell eCDF is close to the red dotted diagonal line suggesting that phase inference is almost equivalent to random phase assignment. For cell group size ≥ 20, the performance is close to optimum. (C) Comparison of PRECISE’s performance for different 3 genes combinations as opposed to using all 4 genes. While all gene sets yielded Δ close to 1 hr, 3 genes allowed phase inference as accurately as 4 genes. Panels A-C correspond to the 23 hr post-Dex synchronisation sample of NIH3T3 cells, where true phase was assigned with respect to *Nr1d1*. (D) Application of SVD to discover circadian phases. Left: Projection of single cells in the eigengene space. Cells belonging to different time points (true circadian phases) are mixed together with no elliptical structure visible. Centre: Projection of samples when gene expression is averaged over 20 cells shows emergence of pure clusters comprising single time points. Right: after averaging over 70 cells, clear and correct clustering emerges along with the expected elliptical structure, allowing for quantitative phase assignment using the coordinates in eigengene space. (E) Comparison in performance of PRECISE and SVD. The median and variance in Δ are systematically lower for PRECISE for the 23 h sample, demonstrating that PRECISE is significantly more accurate than SVD. (F) SVD projection of single cells when 10 simulated genes were added to 6 measured core-clock genes. (G) Purity of SVD clusters derived from single-cell gene expression with increasing simulated gene numbers. All simulated genes had COV corresponding to the least noisy measured core-clock gene, *Nr1d2* (red curve). The blue curve corresponds to simulations where genes had COV 50% less than that of *Nr1d2*.

Like any supervised algorithm, PRECISE makes assumptions regarding the underlying model, noise structure, and additionally requires time-course gene expression data with wavelet based ‘true-phase’ assignments. To confirm that these modeling choices do not qualitatively affect our results, we next explored the performance of unsupervised learning algorithms that do not demand the prerequisites mentioned. Early work demonstrated how Singular Value Decomposition (SVD) can be used to reduce the dimensionality of complex oscillatory gene expression datasets and identify sets of “characteristic modes” or “eigengenes” that represent the global expression profiles [45]. Projection of samples onto the eigengene space leads to emergence of elliptical structures where the angular positions correspond to specific phases of the oscillatory cycle. In order to apply SVD to our test dataset we first generated a gene × cell matrix of dimensions 3 × 250 (taking the same *Bmal1, Nr1d1* and *Tef* combination as shown in Figure 4A-B), whose entries were expression values of 50 randomly chosen single cells from each of the 5 different time points. As shown in Figure 4D (left), projection of the single-cell data onto the top 2 eigengene space failed to cluster cells from the same time point together. We then generated data matrices whose entries were no longer single-cell expression values, but averages over 20 (Figure 4D centre) or 70 (Figure 4D right) cells. 50 such average values per time point were generated by bootstrapping to create matrices of dimensions 3 × 250, and SVD was performed as before. Consistent with the results of PRECISE, we observed that even with 20-cell averages the emergent clusters comprised cells from specific time-points only, and a clear elliptical structure emerged with the correct ordering of the time points at the 70 cell average level (Figure 4D right). We noticed that the samples from the 8 and 32 h time points do not cluster together as might be anticipated. This happens because the circadian clock’s period is not exactly 24 hours, hence they do not represent the same phase. In order to quantitatively compare the performance of SVD relative to PRECISE we inferred the circadian phase of the individual samples by obtaining the *tan*^-1^ of the ratio of their coordinates in the eigengene space and subsequently calculating Δ. We observed that the median and variance of Δ for almost all cell averages were lower for PRECISE in comparison to SVD, suggesting more accurate phase estimation using our supervised learning algorithm (Figure 4E; details in Figure S9 and Section S7).

To check whether the inability to cluster single cells by phase was a result of using just 4 genes, we performed a new experiment where we measured 2 additional core-clock genes, *Cry1* and *Nifl3*, with no improvement in results (Figure S10). We then estimated the coefficient of variation (COV) of the least noisy of these 6 genes (*Nr1d2* with an average COV of 0.81) and computationally simulated 10 additional “clock-like” genes with the same COV, but with different phases (details in Section S7). This total of 16 genes represents approximately the upper limit on the number of core-clock genes that have been suggested to robustly oscillate across tissues. However, single cells remained mixed after SVD dimensionality reduction (Figure 4F), suggesting that it may not be possible to infer circadian phase from single cells using just the core-clock genes. Interestingly, when the number of simulated genes was increased to about 50, clusters with accuracy greater than 90% emerged even at single-cell resolution (Figure 4G). This requirement of 50 genes did not change significantly when the COV was reduced by 50%, to mimic the possible existence of low-noise clock-controlled genes (Figure 4G).

Finally, since previous work has demonstrated the superiority of autoencoders in cyclic phase reconstruction [24], we also checked that an autoencoder applied to the results of SVD makes no difference to our results (details in Section S8 and Figure S11). We also demonstrated the accuracy of across-cell line phase predictions by performing inference on mouse lung fibroblasts (MLG), when PRECISE was trained using time course data from NIH3T3 cells (details in Section S9, Figures S12-S13).

In summary, both the supervised and unsupervised algorithms suggest that true clock phases cannot be recovered at the single-cell level from core-clock genes alone, due to the large intrinsic noise from bursty transcription. However, accurate inference with just ∼ 1 hour deviation is possible with only 3 core-clock genes after averaging over a small group of cells (20-50 cells for PRECISE, ∼ 70 for SVD). Interestingly, our simulations predict that about 50 oscillatory genes with noise levels similar to core-clock genes would be required for accurate phase inference from single cells.

### Non-linear methods fail to cluster single cells by circadian phase using core-clock genes

We next asked if instead of the linear dimensionality reduction technique SVD, non-linear dimensionality reduction methods like UMAP (Uniform Manifold Approximation and Projection) [46] or t-SNE (t-distributed stochastic neighbor embedding) [47] would allow for better separation of the single cells according to their known true circadian phases. Note that unlike PRECISE or SVD, these non-linear methods are not expected to allow for quantitative inference of phase since the elliptical structure (Figure 1A) will not be maintained after projection into lower dimensions. We used both UMAP and t-SNE on our absolute RNA counts to visualize the data in 2D (Figure 5A; details in Methods and Section S10). Annotating the cells based on their known true circadian phases demonstrated that cells from different circadian phases were mixed together with no clear clusters emerging, further emphasizing the fact that inherently noisy circadian gene expression limits correct circadian phase assignment at single-cell resolution. While UMAP and t-SNE are dimensionality reduction techniques, recent efforts have gone towards partitioning scRNA-seq datasets to identify ‘metacells’ representing distinct cell-states, where the variability within a metacell arises from technical rather than biological sources [48, 49]. We therefore asked if cells from within a single time point or circadian phase could be discovered as a metacell and implemented a recently developed algorithm, SEACells, to infer 5 metacells in our dataset. We checked what proportion of cells from different time points were classified into a single metacell [49]. With accurate classification, cells from a single time point would be expected to dominate the population of cells included in a single metacell. However as shown in Figure 5B, all the metacells identified encompassed cells from different time points, suggesting that like UMAP and t-SNE, SEACells also fails to identify the true circadian cell-states as a consequence of the underlying transcriptional noise. Furthermore, the identified cell-states were not robust – the 3 methods grouped very different sets of single cells into clusters, which further depended on the specific parameter choices (Figure S14, Methods and Section S10).

**Figure 5:**
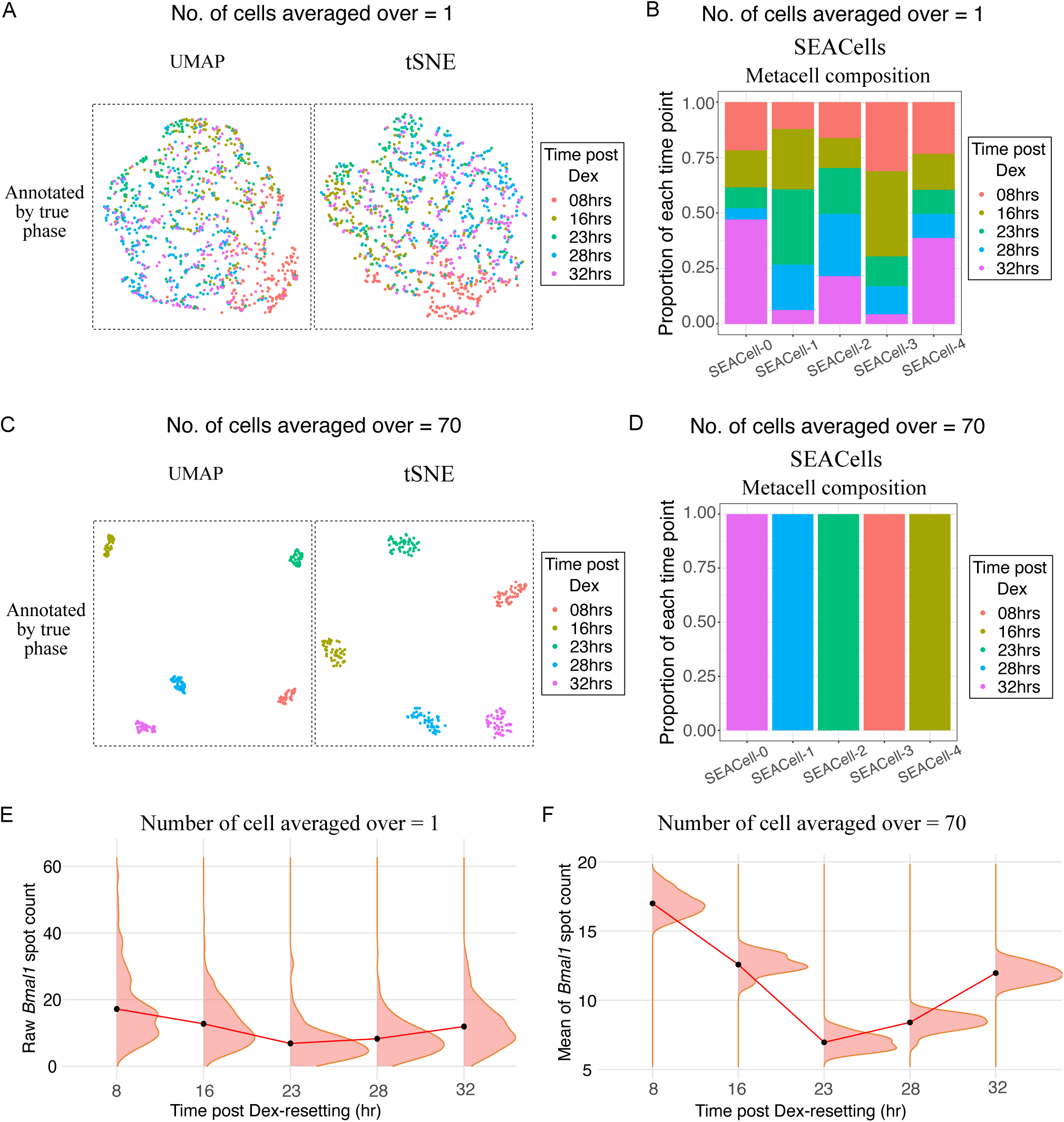
Transcriptional noise prevents meaningful clustering of single cells by cell-state discovery algorithms. (A) All four genes *Bmal1, Nr1d1, Nr1d2* and *Tef* were used to obtain UMAP and t-SNE based dimensionality reduction. Cells annotated by the known true circadian phases (times post Dex resetting) in the UMAP and t-SNE projections suggests that cells from the different circadian phases were mixed together with no clear emergent clusters. (B) SEACells based identification of ‘metacells’ for our single-cell circadian clock dataset. Metacell composition shows the proportion of cells from different true circadian-states that are included in each metacell. Similar to panel A, each SEACell identified metacell comprises cells from different true states. (C-D) Application of UMAP, t-SNE and SEACells on a dataset averaged over 70 cells successfully identifies pure clusters, with each cluster comprising cells from only one particular true circadian state. (E) Distributions of raw spot counts for the gene *Bmal1* in single cells across the 5 test time points post Dex-resetting. While the means of the distributions (black dots) were shifted, the distributions were highly overlapping, thereby limiting cell-state inference at single-cell resolution. (F) Distributions of mean *Bmal1* spot counts across time post Dex, when gene expression was averaged over groups of 70 cells. The distributions of the means are clearly well separated, allowing accurate cell-state discovery.

Since the results in the previous section had demonstrated our ability to accomplish accurate phase inference only after averaging, we next tried the same strategy on all the 3 methods. We generated 50 samples per time point, where each sample was obtained by averaging gene expression over 70 randomly chosen cells from within that time point. We put together all these samples from 5 time points and explored the performance of each of these algorithms on the averaged dataset. Consistent with the results of our last section, we now observed that UMAP and t-SNE projections in 2D clearly separated out the different time points (Figure 5C). The same was true for SEACells – each metacell identified now comprised samples from only a single time point, as shown in Figure 5D. Therefore irrespective of the cell-state discovery algorithm we used, we could recover the true circadian states only after averaging.

### Biological, not sampling noise prevents correct single-cell phase assignments

To understand what enabled accurate phase assignment for cell groups but not single cells while using the core-clock genes, we visualized the distribution of raw spot counts for the genes across the various time points (the distributions for *Bmal1* are shown in Figure 5E, the rest are shown in Figure S15). While the means of the distributions (black dots in Figure 5E) were shifted as expected from our analysis in Figure 1F, the distributions were strongly overlapping for the different circadian phases. However, the distribution of the means were well separated since the variance of each distribution was much reduced (Figure 5F). Note that though the means of the 16 and 32 h time points are the same (Figure 5F), the presence of the other genes will allow for unique state assignments. Dimensionality reduction and cell-state discovery algorithms, with their inherent approach of clustering cells with statistically similar expression profiles, can therefore discover the correct phases only after averaging.

Could sampling noise have contributed to the non-identifiability of single-cell circadian phases? In sequencing based methods like scRNA-seq, only a small fraction of the total cellular RNA (upto ∼ 30% with modern protocols [50]) can be captured, leading to sampling artefacts. However, single-molecule imaging based methods like smFISH largely overcome this issue. In our SABER-FISH setup, the high colocalization fraction of 90-95% (see Figure S1 and Section S1) combined with simple probability arguments demonstrate that our probes capture around 95% of the total transcripts (see Section S11 for details). To further check if the 5% of missed transcripts could potentially lead to issues of circadian phase identification, we ran a number of simulations. We generated synthetic distributions of single-cell RNA counts for four different genes, representing two distinct circadian phases. We utilized the NB parameters derived from fits to our training datasets, resulting in the underlying distributions being strongly overlapping. For each of the two phases, we sampled 200 or 1000 individual cells each characterized by a vector of absolute counts of 4 genes sampled from the underlying distribution. The counts per cell thus generated are the ‘true counts’ or the actual RNA numbers in different cells. From these, we ultimately obtained the ‘observed counts’ by sampling from the true counts with some probability *<*1. We applied UMAP to the observed counts matrix of RNA per cell and annotated the samples by their true phases to analyze their clustering patterns. As shown in Figure S16, regardless of whether the sampling probability was low (0.20-0.30; characteristic of scRNA-seq) or high (0.90-0.95; characteristic of smFISH), when the underlying distributions were overlapping (as is the case with the circadian clock), phase inference was possible only after averaging.

Overall, this section demonstrates that our experimental observations on the circadian clock phases (correct clusters emerge only upon averaging the core-clock genes) arises when biological noise drives the gene expression distributions of different phases to strongly overlap with one another. In such cases, irrespective of the level of sampling noise, single-cell phase assignments will not reflect the true circadian phases.

### Averaging over locally proximal cells allows spatially-resolved circadian phase inference

In the last sections we demonstrated that single-cell circadian phase inference is fundamentally limited by the transcriptional noise associated with core-clock gene expression. However, averaging over small cellular groups from within the same phase leads to reduction of this noise, thus allowing accurate phase estimation for synchronised populations of cells. A key question that therefore arises is regarding the selection of cells to be averaged over, in order to accurately represent the underlying circadian phase. For a synchronized population of cells where the underlying assumption is that all cells are in the same phase, a random choice of cells is clearly appropriate. When we performed phase inference for 50 samples, each of which was derived from gene expression averaged over 20 randomly chosen cells from our Dex-synchronised test dataset, we observed that the inferred phases were localised to specific regions on the circle (Figure 6A). To quantitatively measure the phase coherence, we computed the Rayleigh statistic, whose value is close to 1 when the data is localised on a circle and 0 when the data is uniformly distributed (see details in Section S12). The high Rayleigh statistics observed for the synchronized population of cells (Figure 6A) as well as the clear separation of clusters seen earlier (Figures 5C-D and 4D) demonstrate that random selection of cells is sufficient.

**Figure 6:**
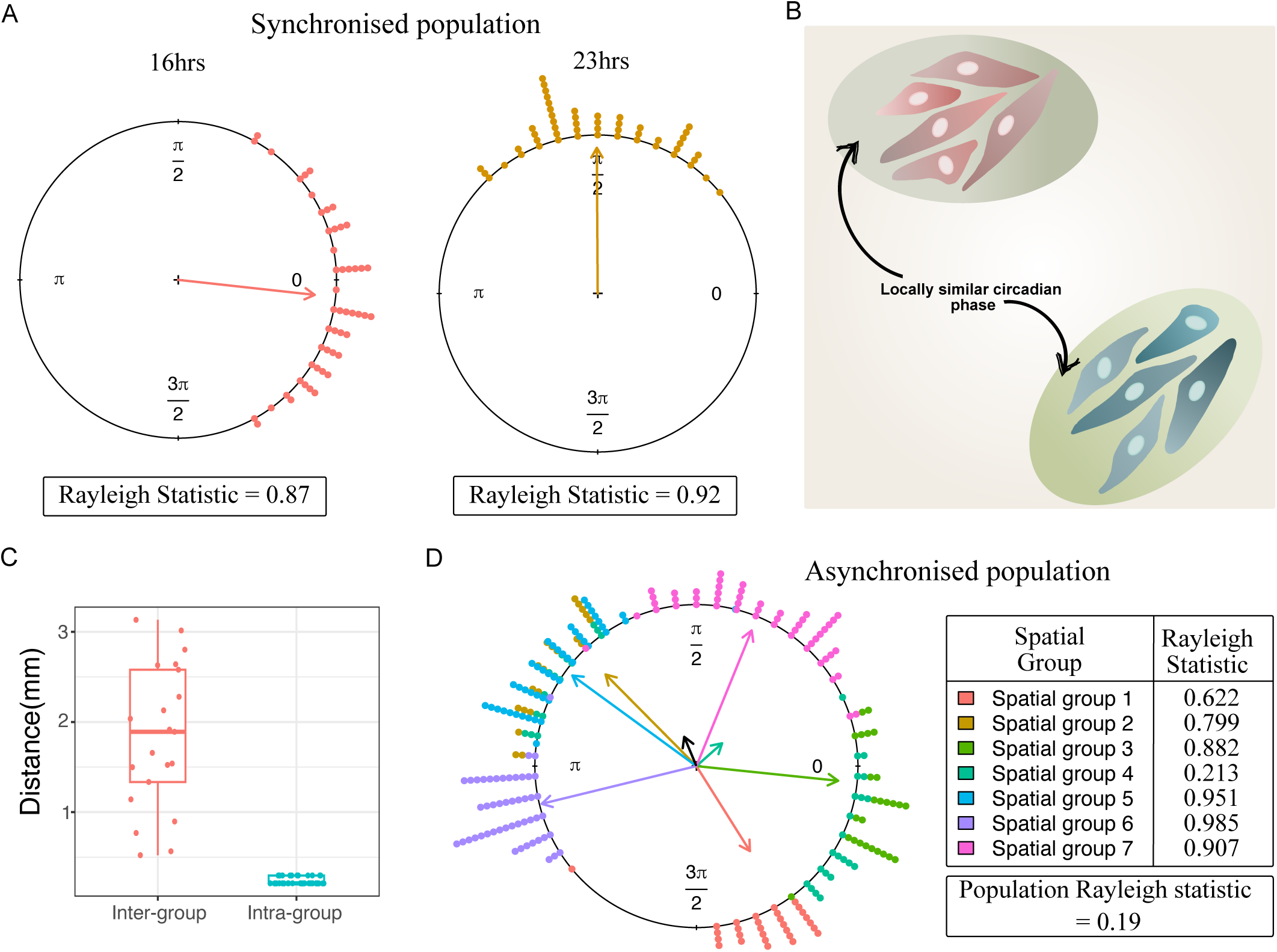
Averaging over locally proximal cells allows spatially-resolved circadian phase inference. (A) In synchronized populations, circadian phase was assigned 50 times (solid circles) via averaging over 20 randomly selected cells on each occasion (left-16 h and right-23 h time points). The Rayleigh statistic quantifying the spread in the inferred phases is close to 1 in both cases, suggesting strong phase coherence. (B) For an asynchronous population, local regions of cells are expected to have similar circadian phases due to paracrine signaling or lineage effects, but distinct phases will be maintained for spatially distant cell groups. (C) Comparison of the distances within and across 7 spatial groups of cells that were imaged in an asynchronous population of NIH3T3 cells. The distances of cells within each group (intra) were distinctly less than the distances across groups (inter). (D) Circadian phases inferred for different spatial groups of cells (denoted by different colours) in the asynchronized population. Similar to panel A, each spatial group comprised 50 phase measurements (solid circles) after averaging over 20 cells randomly chosen from within that group. The overall Rayleigh statistic is close to 0 (black arrow), demonstrating a near-uniform phase distribution across space. However, high phase coherence is observed within the different spatial groups as quantified by the high individual Rayleigh statistics (coloured arrows).

On the other hand, how should one select cells for averaging from an asynchronized population? This is typically the case for most biological systems (for example developmental cell-states), which unlike the circadian clock cannot be well synchronized or are not often found in synchronized states. For such asynchronous populations comprising cells from various states or phases, spatially proximal cells are likely to be representative of the same cell-state. In the context of the circadian clock, a recent study suggested that paracrine signaling via TGF-*β* causes local regions of cells to have similar circadian phases but spatially distant cells to have distinct phases (Figure 6B) [51]. Furthermore, if phases are decoded correctly, a uniform circadian phase distribution is expected to emerge from an asynchronous population, as has previously been demonstrated using live-cell imaging [52]. We utilized these two ideas to see if spatially resolved phase inference could be performed reliably. We carried out SABER-FISH on an asynchronous population of NIH3T3 cells (details in Section S12), followed by phase inference using PRECISE on spatially averaged gene expression values. In total we had 7 separate spatial groups in our data, and we confirmed that these groups were sufficiently separated from each other as indicated by larger inter-group distances compared to the intra-group distances (Figure 6C and Figure S17). For the cells belonging to a particular spatial group, we performed 50 instances of phase prediction, where for each instance gene expression averaged over 20 randomly selected cells was used as input. We executed similar phase inference for each of the remaining spatial groups present in our data and visualised the inferred average phase (*ϕ_pred_*) on a circle. As shown in Figures 6D and S18, both the expectations were met – (i) a uniform distribution of inferred phases was found for the entire population (black arrow corresponding to the population Rayleigh statistic) and (ii) high phase coherence was observed within each spatial group as characterised by the high individual Rayleigh statistics (colored arrows). Together, these results suggest that choosing spatially proximal cells may be good strategy to perform the necessary averaging of gene-expression before decoding circadian phases.

To summarize, averaging over random groups of cells is sufficient to detect the true circadian phases only in synchronized cell populations. However, when dealing with an asynchronized population that is expected to have cells in a mixture of phases, we suggest a simple strategy of averaging over spatially proximal cells to recover the true underlying phases. Our results provide a principled approach towards decoding circadian phase in a spatially resolved manner.

## Discussion

Here we undertake the first systematic study of the limits of measuring circadian phase from a single snapshot of core-clock RNA levels, in terms of the minimum number of cells and genes required for accurate phase inference. To circumvent the significant technical limitations of single-cell sequencing technologies that make it challenging to answer questions on fundamental limits, we utilize a highly sensitive and accurate smFISH technique called SABER-FISH [40] to measure RNA levels in single cells. Intriguingly, we find that the core-clock gene RNA ratios that are widely used to define circadian phase in population measurements, cannot be directly used to ascribe meaningful circadian phases to single cells. We carefully demonstrate that this limitation arises due to biological, not technical noise – the bursty nature of transcription results in single cells from different circadian phases having large overlap in core-clock RNA levels, even when multiple clock genes are measured. Remarkably however, averaging RNA levels over small groups of 20-70 cells allows accurate phase inference with less than 1 hour error, using just 3 core-clock genes. 20-70 cells is small enough to provide near single-cell resolution, but large enough to average out the effects of transcriptional noise. We believe that this result will provide a guideline for principled approaches to high resolution clock phase detection studies in the future. Indeed, we also demonstrate how an approach of local averaging can be used to detect spatially heterogeneous circadian phases in an asynchronous population of cells. This opens up the exciting future possibility of spatially resolved endogenous circadian phase decoding in solid tissues; if achievable, this could allow resolving the origins of clock dampening in various disease contexts. Interestingly, increasing the number of oscillatory genes computationally to about 50 allowed for accurate clustering by true circadian phase even at single-cell level. However, given that at maximum ∼10-20 core-clock genes are known to consistently and robustly oscillate across tissues, it is unlikely that 30-40 additional genes with similar (or less) noise levels compared to the core-clock genes can be found in any tissue, to enable single-cell phase inference. Further studies will be required to ascertain the validity of this prediction.

Our study has a number of limitations that will need to be clarified in future work. First and foremost, all our results have been derived from cell lines. While transcriptional bursts have been reported in tissues [53], there might be additional noise reduction mechanisms in the *in vivo* context that may change the level of coarse-graining or gene numbers predicted by our current results. Second, Dex resetting might be imperfect in synchronizing single-cell circadian clocks, leading to a mixture of true states even within a single time-point post Dex addition. While possibly existent to some degree, this limitation is unlikely to affect our results significantly. Since Dex is a well established strong resetting agent generating ‘type-0’ phase-transition curves, most single cells are expected to be in the same state upon Dex treatment [52]. If there were a number of other states mixed in due to incomplete Dex action, one would have expected one large cluster along with some small clusters upon phase inference. We do not see this behavior – single-cell phase inferences are uniformly distributed (Figure 4A) and clusters are also well mixed in the true states (Figure 5A-B), ruling out any major effects of incomplete Dex synchronization. Finally, it remains possible that more powerful computational algorithms developed in the future could push down the limits we discover in this study. For example, rapid developments in representation learning methods, or methods using RNA velocity could potentially be leveraged to challenge the limits identified here.

The prediction of this study that inferring circadian phase is not possible from single cells using just the core-clock RNA levels, although feasible using population measurements, may appear surprising at first. However, it is increasingly becoming clear that functional cell states in general need not emerge from RNA levels alone, but likely from a combination of other cellular components like protein levels, the epigenome and chromatin architecture [54]. With less associated noise due to higher copy numbers, measurement of only a few protein levels could conceivably allow for single-cell level determination of circadian phase. However, if only RNA levels can be experimentally accessed, coarse graining may become necessary, as our work suggests. Interestingly, we found a conceptually similar result in a different context, in an older study using scRNA-seq with careful spike-in controls to distinguish biological versus technical noise [55]. This study demonstrated that to recover the same transcriptome and number of measurable genes as obtained from bulk RNA-seq measurements, RNA levels from groups of 30-100 cells needed to be averaged over [55]. This number is remarkably similar to our finding of 20-70 cells, though in a very different context. These results together suggest a potentially general principle – even if technical noise can be minimized, averaging over a small group of cells as opposed to using single-cell RNA levels might allow more robust and meaningful assignment of cell-states in a variety of biological contexts, beyond the circadian clock.

## Author contributions

S.C. conceptualized and designed the study with help from A.N., N.V. established and optimized the SABER-FISH protocol, T.M. developed the multiplexed version of SABER-FISH, collected training datasets with A.N. and performed initial data analysis, N.V. and T.M developed the initial image analysis pipelines, A.N. collected all test datasets across cell lines, built the final image analysis and registration pipelines, developed the supervised algorithm PRECISE and performed the computational analyses, A.N. and S.C. wrote the paper with inputs from N.V. and T.M., S.C. supervised the work and raised funding for the project.

## Acknowledgments

S.C. acknowledges funding from SERB (Government of India) under project number SPR/2021/000486 as well as intramural funds from National Center for Biological Sciences–Tata Institute of Fundamental Research (NCBS-TIFR) via Department of Atomic Energy, Government of India, Project Identification No. RTI 4006.

## Methods

### Cell culture

NIH3T3 (mouse; ATCC, CRL-1658) and MLG (mouse; ATCC, CCL-206) were grown in DMEM (Genetix, cat. no. CC3004.05L) supplemented with 10% (vol/vol) serum (Himedia, cat. no. RM10432-500ML) and Penicillin - Streptomycin (Gibco, cat. no. 15140122). All cells were incubated at 37*^◦^*C and 5% CO_2_. In all experiments, cells were grown to about 70 − 80% confluency before fixation and imaging.

### SABER-FISH for multiplexed RNA imaging

To visualize endogenous RNA of different clock genes that are produced in low numbers, we utilised a recently developed technique Signal Amplification by Exchange Reaction - Fluorescence *in-situ* Hybridisation (SABER-FISH) [40], with all reagent concentrations and reaction times taken from the updated protocols provided in [41]. To generate the training dataset we performed multiplexed imaging of 4 clock genes – *Tef, Bmal1, Nr1d1* and *Nr1d2* for fixed monolayers of cells over every 4 hours for 48 hours after Dexamethasone treatment. To generate NIH3T3 cell line test dataset we obtained the counts of the same four clock genes over time points – 8, 16, 23, 28 and 32 hrs post Dex-resetting. To generate MLG cell line test datasets we measured the 4 clock genes included in training data across time points – 8,16, 23, 28 hrs post Dex-resetting. In a separate experiment performed on the NIH3T3 cell line, we obtained the absolute RNA counts of 6 clock genes – *Tef, Bmal1, Nr1d1, Cry1, Nifl3* and *Nr1d2* over 6 different time points post Dex-resetting – 8, 12, 16, 20, 24 and 28 hrs. For each gene, about 50 primary hybridization probes were designed to tile the gene, and extended to various lengths using the Primer Exchange Reaction. These extended probes (or concatemers) were hybridized overnight, following which imager sequences attached to fluorophores were hybridized onto the concatemers. The cells were then imaged on a Nikon Ti2-E epifluorescence microscope with a 60x oil objective, using a Hamamatsu sCMOS camera (ORCA FLASH 4.0). Multiple rounds of imaging were performed by using Formamide washes, which removes the secondary imagers but not the primary probes. A cytoplasmic stain CellMask (Invitrogen, cat. no. H32720) was also included in the different imaging rounds to allow segmentation followed by spot counting per single cell. More details of the experimental protocols and standardisation procedures are provided in Section S1.

### Image analysis to quantify single-cell gene expression

Multiplexed SABER-FISH images were processed using custom image analysis pipeline based on ImageJ and Cellpose [56, 57]. We detected and extracted the spot coordinates for the different genes from maximum Z-projected images using the RS-FISH plugin in ImageJ [58]. Segmentation of the cells was performed based on contrast-enhanced CellMask images using Cellpose. We wrote custom python-based scripts to perform image registration to identify the same cells across the different rounds of images, and assign spot counts to each individual cell. Further details and parameter values used in the various algorithms are included in Section S2.

### Bulk gene expression measurements using qPCR

Quantitative real-time PCR (qRT-PCR) assay was used to obtain bulk gene expression levels for the different clock genes. We utilised Dexamethasone-resetting to synchronise cells, followed by Trizol based extraction of total cellular RNA using the Purelink RNA extraction kit (Invitrogen, cat. no. 12183018A). cDNA conversion of the cellular RNA was performed using SuperScript III First-Strand Synthesis System (Invitrogen, cat. no. 18080051). qRT-PCR was performed in 10*µ*L final reaction volume, with three replicates per sample, using iTaq Universal Sybr-Green supermix (Biorad, cat. no. 1725121). We performed qRT-PCR for 2 clock genes – *Bmal1* and *Nr1d2*, and the constitutively expressed gene *Gapdh*. ΔΔCt analysis was performed to obtain the clock genes’ expression values as fold change with respect to *Gapdh*. Further details related to the experiment and analysis are included in Section S4.

### PRECISE: Predicting Circadian phase from Stochastic gene Expression

We developed a new algorithm, PRECISE (**PRE**dicting **CI**rcadian phase from **S**tochastic gene **E**xpression), to infer the circadian phase of a test sample from RNA measurements in a supervised manner. The input training data for PRECISE comprises the normalized mean RNA levels as functions of circular phase values (obtained from time post Dexamethasone synchronisation expression data), as well as the errors of the normalized means. At its core, PRECISE uses Gaussian Processes (GPs) with a non-stationary kernel to learn the oscillatory patterns of the different genes from the input training data. PRECISE builds on ODeGP [43], an accurate and sensitive oscillation detection algorithm that we previously developed, to learn the posterior means and variances of the time series of each gene. The power of this approach lies in the fact that GPs provide a non-parametric inference framework in function-space, and provide analytic expressions for the posterior mean and variance (for a basic introduction to GPs and its application in oscillation detection, see [43].) These expressions for the posterior means and variances are subsequently used to construct a likelihood function that allows calculation of the likelihood of any particular phase given gene expression measurements of a test sample. The likelihood curve tends to have multiple peaks since our training data spans 48 hours post Dex-synchronization. Hence the final predicted circadian phase assigned to the sample, *ϕ_pred_*, is calculated by taking the circular mean of the phases corresponding to the likelihood peaks. Mathematical details of PRECISE and its implementation are provided in Section S6.

### Assignment of ‘true’ circadian phase to time-points post Dex-resetting

The time-points post Dex resetting were converted to phase angles using the procedure broadly described below (for details see Figure S5 and Section S6). The true or global phase of the training or test samples can be determined with respect to any one (arbitrarily chosen) clock gene. For instance, a peak of *Bmal1* could be arbitrarily assigned phase 0 and the next peak as phase 2*π*. For the training samples, we first bootstrapped the data for a particular gene from each time point to generate mean expression and errors of the mean as functions of time, as described in Section S6. The bootstrapped data is then subjected to normalization by which the mean and standard deviation over time was made 0 and 1 respectively as described in Section S6. This normalized input was utilised by ODeGP [43] to determine the posterior mean fit to the data, allowing extraction of the normalized expression values at any arbitrary time point, beyond the experimentally measured ones. Normalized expression was then extracted at discrete time points with a given sampling rate from the posterior mean, and provided as input to pyBOAT [44] for determining phase values. pyBOAT is an algorithm that uses wavelet-transforms to determine the phase of the different samples as functions of time [44]. The phases corresponding to the time points present in our experiments were then extracted and used as phase estimates of the experimental time-points. The phase estimates of the 13 time points used as training data are denoted *ϕ_i_* (*i* = 1, 2*, …,* 13) while the phase estimates of the test samples are denoted *ϕ_true_*. More details and parameter values of softwares used are provided in Figure S5 and Section S6.

### Calculation of deviation of inferred phase from true circadian phase

Once PRECISE assigns a predicted phase *ϕ_pred_* to a test sample, its deviation from the true phase *ϕ_true_* was calculated in two steps as follows (see Section S6 for details):

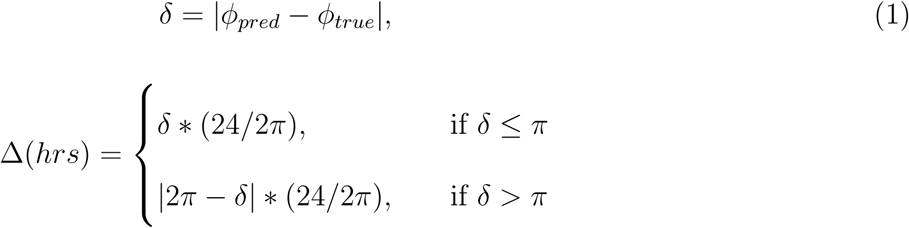

Thus the deviation in predicted phase, Δ, always lies between 0 to 12 h.

### Unsupervised dimensionality reduction and cell-state discovery algorithms

We checked whether the results of a variety of unsupervised cell-state discovery algorithms are consistent with PRECISE. To this end, we used SVD (Section S7), lower-rank approximation of the data using SVD followed by an autoencoder with a circular bottleneck layer (Section S8), UMAP, t-SNE and SEACells (Section S10) to discover circadian clock states. Details of each method are provided in the Supplementary Information as indicated.

