## Supplementary Information for "Transcriptional noise sets fundamental limits to decoding circadian clock phase from single-cell RNA snapshots"

**S8 Neural network based circadian phase inference. 36**

**S9 Circadian phase prediction across different cell lines. 38**

**S10 Cell-state discovery algorithms 39**

### S1 Quantifying circadian clock gene-expression using SABER-FISH

#### 1. Cell culture reagents and protocols.

NIH3T3 (mouse; ATCC, CRL-1658) and MLG (mouse; ATCC, CCL-206) cells were grown in DMEM (Genetix, cat. no. CC3004.05L) supplemented with 10% (vol/vol) serum (Himedia, cat. no. RM10432-500ML) and Penicillin - Streptomycin (Gibco, cat. no. 15140122). All cells were incubated at 37°C and 5% $CO_2$. A passage 15 (p15) NIH3T3 (Mouse Embryonic Fibroblast) line was used to generate the training data while p9 NIH3T3 line and p70 MLG (Mouse Lung Fibroblast) line were used to collect the test data points.

#### 2. Details of SABER-FISH gene expression measurements.

##### 2. .1 Overview of SABER-FISH.

To image multiple genes in the same single cells, we used a recently developed technique, Signal Amplification by Exchange Reaction (SABER) - FISH [1]. In this technique, many primary probes complementary to a single RNA are designed to tile the RNA of interest. These probes are linked to primers with a specific repeat sequence, and first subjected to a 'Primer Exchange Reaction' (PER) *in-vitro* [1, 2]. This reaction extends the repeat sequences to create overhanging 'concatemers', which then act as platforms for binding multiple secondary imager probes labeled with fluorochrome (main text Figure 1B). This process thus leads to amplification of the signal coming from a single RNA molecule. This technique also allows for easy multiplexing to rapidly image many genes in the same single cells. Multiplexing is achieved by using unique repeat sequences for the concatemers and corresponding complementary imager sequences, for the different RNAs of interest. Primary probes for all RNAs are hybridized in one overnight step, followed by imaging of groups of genes across different imaging rounds. Each imaging round comprises imager hybridization and microscopy, followed by Formamide washes to selectively remove the imagers while keeping the primary probes intact for the next round of imager hybridization and imaging.

#### 2. .2 **Validation of SABER-FISH using a colocalization assay.**

In order to standardise and validate the SABER-FISH protocol for our cell lines, we designed probes targeting the *Cbx5* gene. *Cbx5* is a highly expressed gene that has been well studied and characterized before using smFISH [1]. The primary probes were divided into two separate groups (purchased as IDT oligo-pools), where the difference was the concatemer sequence linked to them. The imager probes complementary to the two different concatemers were bound to two different fluorescent dyes – Atto488 and Atto545. We proceeded to perform the colocalization assay on NIH3T3 cells. We extended the two oligo-pools using PER (reaction times and reagent concentrations were as per [2]). After fixation of the cells using 4% Paraformaldehyde (PFA) for 15 mins at room temperature, we allowed primary hybridisation of both the extended probes simultaneously overnight at a specific temperature. Secondary hybridisation of both sets of imaging probes was performed the following day. If the specificity of the assay was high, we expected spots in the two different channels to co-localize. We obtained approximately 95% colocalization between the spots in the two channels when primary hybridisation was performed at 45°C.

In order to quantify the extent of colocalization between the spots present in the two different channels, we assembled a custom spot counting pipeline. The steps of the pipeline are briefly mentioned below:

1. First we extracted the coordinates of the smFISH spots in both the channels. We obtained the maximum Z-stack projection for the images, followed by RS-FISH [3] based spot detecting and counting. We manually set the thresholds (separately for both channels) to obtain the coordinates which were saved as separate .csv files.

2. After obtaining the coordinate files, using a custom python-based code we obtained the percentage of spots colocalizing in our images. This was done by considering a fixed spot in the first channel and performing a search for spots in the second channel within a 4-6 pixel radius. If we detected a spot in the second channel within the given radius, we considered those spots as colocalised. This procedure was repeated for all the spots within the first channel and then the percentage of spots that colocalise with a spot in the second channel was computed.

Following the above procedure we performed the colocalization assay on *Cbx5* spots and quantified the percentage of colocalization. We performed this on both cell lines used in this paper, NIH3T3 and MLG. As visually evident from Figure S1, when we merged the images obtained in the green and red channels, the majority of spots appeared yellow suggesting high colocalization. This proves the sensitivity of the assay and that $\sim$ 95-96 % of the spots are real signal, not noise.

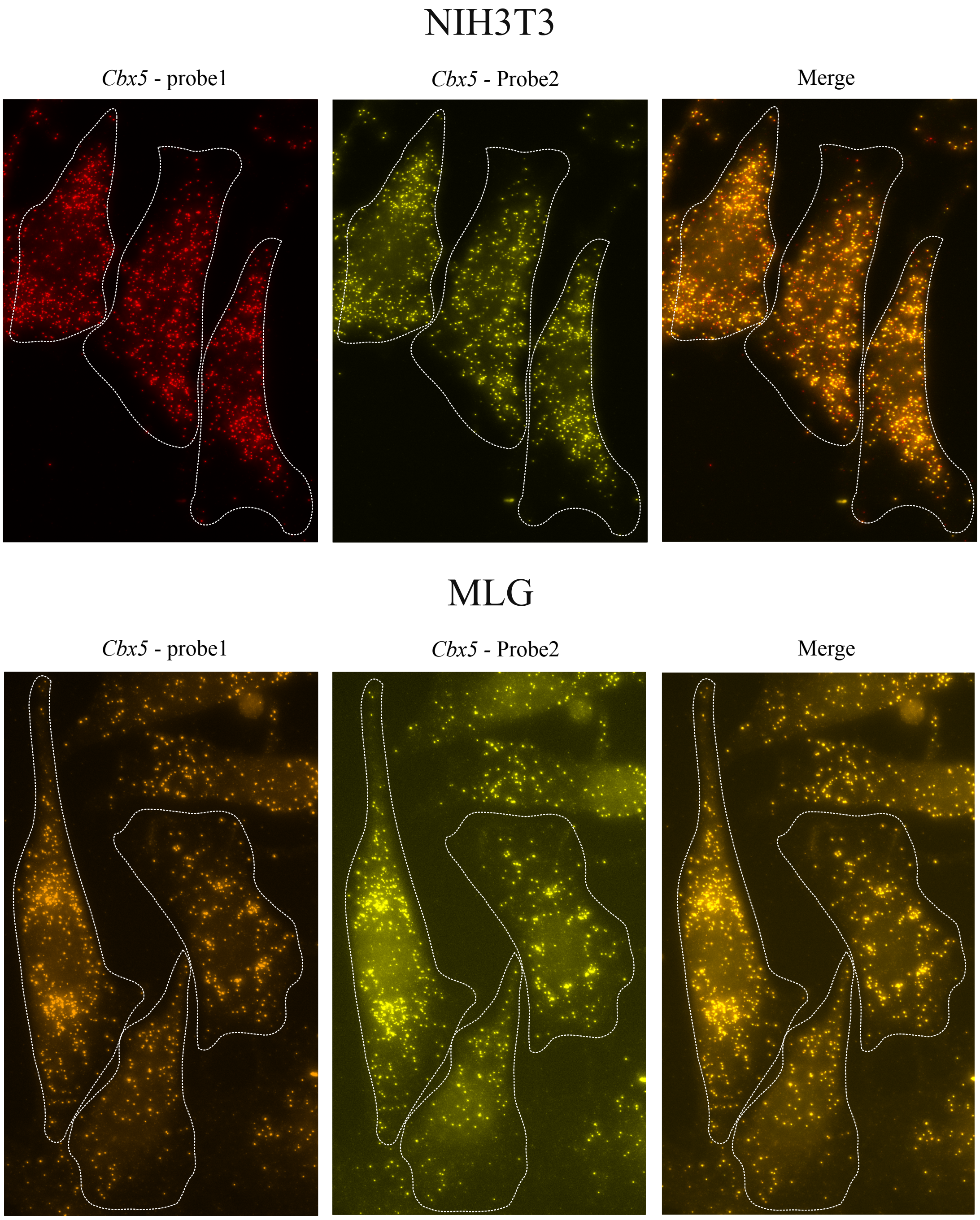

Figure S1: Colocalization assay to validate SABER-FISH.

#### 3. Dexamethasone-synchronisation of NIH3T3 cells to generate training datasets.

To generate cells in well defined and distinct circadian phases, we used Dexamethasone (Dex) to synchronize the cellular clocks in culture. Dex has been shown in many earlier studies to be a strong resetting agent that shifts the clocks of cells to the same phase, irrespective of the initial phase of the cells. Additionally, it is known from ours and other studies that differences in cell density can affect inter-cellular coupling and the synchronization of clocks, hence we designed a protocol to control for cell density while collecting cells at different times post Dex-synchronization. This protocol was used to generate the training data for NIH3T3 cells, and is shown in Figure S2. In brief, instead of Dex addition and cell-fixation at different time points, we added Dex at different time points and collected the cells at one time. This allowed for maintaining roughly equal cell numbers at time of fixation. The protocol was divided into two parts for ease of implementation – time-points T6, T10, T14, T18, T22 were collected at the same time while time-points T26, T30, T34, T38, T42, T46, T50, T54 were collected at a different time (Figure S2). These two sets were initially seeded with different seeding densities, to control for the number of days in culture, to approximately get similar cell numbers across all time points at time of fixation.

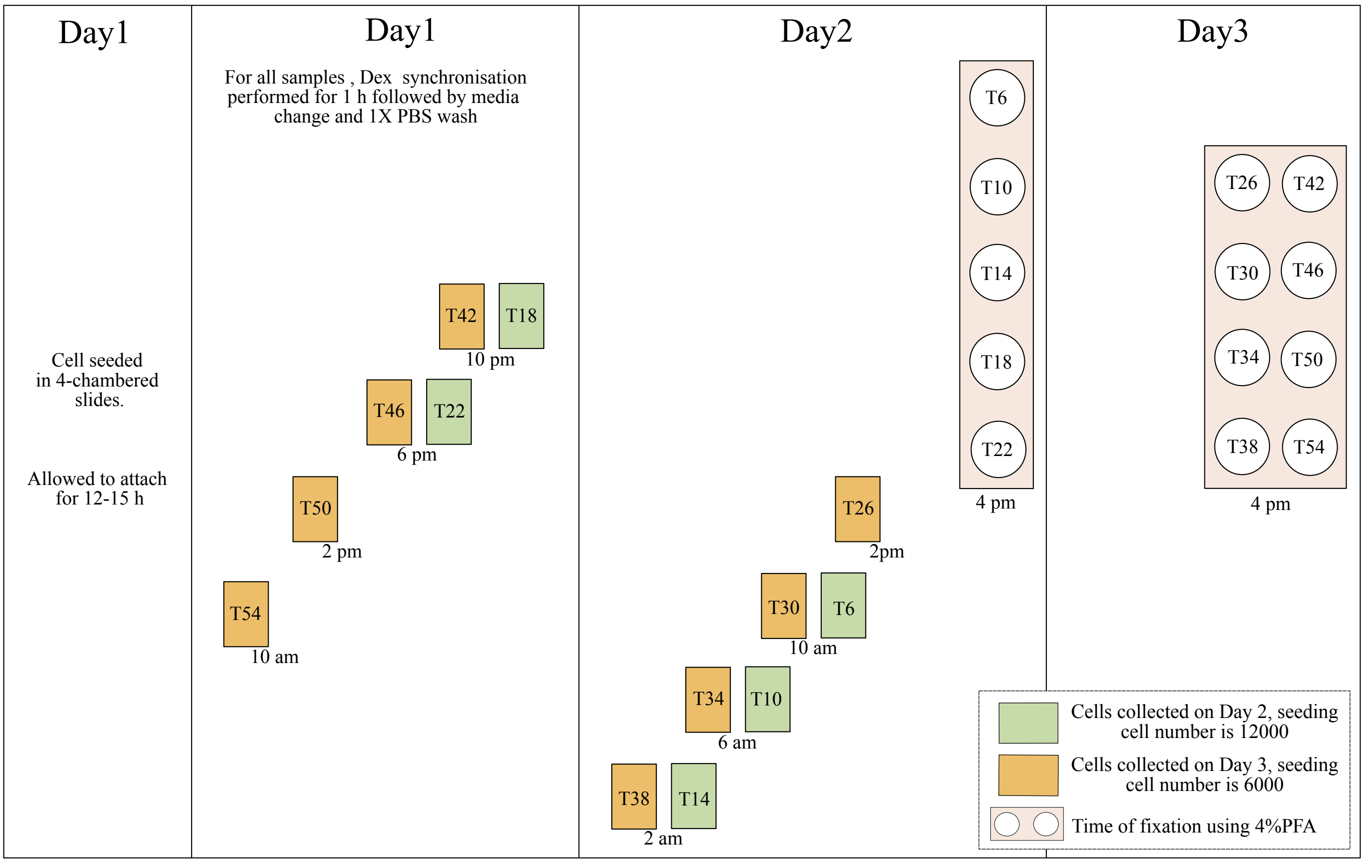

Figure S2: An overview of synchronisation schedule used to generate the training data

At the beginning the cells were seeded into multiple 8-chambered slides (Nunc Lab-Tek II slides, Catalog no. 155409) where 2 wells corresponded to a specific time point. We collected data over 48 h (two circadian cycles), while maintaining 4 h intervals between consecutive time points. Therefore, we seeded 26 wells across four 8-chambered slides corresponding to the 13 different time points under consideration. In order to eliminate the initial transients of Dexamethasone treatment, we refrained from collecting data in the initial 6 h post synchronisation hence our time points ranged between 6 to 54 h. For Day 2 set of samples, 12000 cells were seeded per well and for Day 3 set of samples 6000 cell per well were seeded. We allowed the cells to attach for ~12-15 h post seeding, then began the synchronisation of the cells using Dexamethasone. For synchronisation, we incubated the cells for 1 hour in complete media containing Dex at a final concentration of 100nM. After 1 hour we removed the Dex and added fresh complete media after a single 1X PBS wash and recorded the time as T = 0. Mentioned in Figure S2 are times corresponding to the T = 0 for the different samples. Since the doubling time of NIH3T3 cell line is ~24 h, hence this protocol allowed for about 2 divisions for Day 2 samples and ~ 3 divisions in case of Day 3 samples, leading to all wells having approximately same density at time of fixation. At the time of fixation, we removed the media from all the wells, followed by a single 1X PBS wash, and proceeded to incubate the wells at room temperature in 4% PFA for 15 mins.

#### 4. Quantifying single-cell clock gene expression using SABER-FISH.

We performed the Primer Exchange Reaction (PER) as described above and in [1] to separately extend the oligo-pools containing DNA sequences complementary to the four clock genes *Bmal1, Nr1d1, Nr1d2* and *Tef.* Post fixation we performed primary hybridization of the extended oligo-pools for all the clock genes simultaneously at 45°C overnight. Secondary hybridization was then performed for two clock genes, allowing for parallel imaging of those two genes. We used the cytoplasmic marker CellMask (in the DAPI channel) to stain the cytoplasm of the different cells to allow for segmentation post-imaging. We imaged ∼200-250 cells per time point sample (considering both wells that correspond to one time point), and preserved the coordinates of the Fields of View (FOVs) for the different samples which later allowed us to image the same cells again to obtain counts data for other clock genes. We then removed the imager sequences using 60% Formamide wash and hybridized the imager sequences for the next two genes, thus allowing simultaneous measurement of absolute RNA counts of 4 clock genes at level of single cells across different time points over 48 h.

#### 5. Generating test datasets for the NIH3T3 and MLG cell lines.

Following the above protocol we also generated test samples by synchronising cells over a few different test time points. For the NIH3T3 cell line test data we collected samples corresponding to 8, 16, 23, 28 and 32 h post Dexamethasone treatment. We were able to image ∼200-250 cells for this cell line. While for MLG cell line, we had time points 8, 16, 23 and 28 h post Dexamethasone samples and imaged ∼150-200 cells per time point. We used 4-chambered slides (Nunc Lab-Tek II slides, Catalog no. 155382) for the test time points and performed synchronisation continously throughtout 32 h hence the seeding density was the same for all wells both in case of NIH3T3 (30000 cells per well) and MLG (30000 cells per well).

In a separate experiment performed on the NIH3T3 cell line (p14), we collected samples corresponding to 8, 12, 16, 20, 24 and 28 h post Dexamethasone treatment. We were able to image 220-300 cells for this cell line and measured the expression value for 6 clock genes namely – *Bmal1, Nr1d1, Nr1d2, Tef, Cry1* and *Nifl3* at single-cell resolution.

### S2 Image analysis to obtain single-cell spot counts of different clock genes.

In order to extract the single-cell spot counts of different clock genes, we assembled a custom image analysis pipeline that utilised the softwares: ImageJ and Cellpose along with python based codes. The steps followed to process the images from a single time point across the two rounds of smFISH hybridisation are as follows:

1. First we extracted the coordinates of the smFISH spots in both channels for each round of hybridisation. We wrote an ImageJ Macro to batch process the images from each round separately. We first obtained maximum Z-stack projection for the yellow and far-red channels separately, followed by RS-FISH plugin based spot detection by using manually set thresholds (separately for each channel) to obtain the coordinates of the spots in the channels. The coordinates for each channel was saved separately as .csv files. The maximum Z-stack projection in the blue channel which is the CellMask cytoplasmic stain was subjected to histogram equalization to enhance contrast which enabled us to perform segmentation of single cells. These were saved as .tiff files. This was repeated for the second round of images as well.

2. We next trained the Cellpose software using the contrast-enhanced cellmask images for some FOVs (Field of View) of the time point to obtain the segmentation masks as png files and used python-based code to batch process the rest of the images. Thus we had two masks per FOV (corresponding to the two smFISH rounds) at the end of this step. Cellpose is designed such that in each of the segmentation maps all pixel values that correspond to a particular cell are assigned a specific numeric value which we treat as a "unique ID" for the cell. This numeric value for the same cell across the two masks (corresponding to the two smFISH rounds) may be different. Thus, we needed to apply image registration to the two masks in order identify the same cell across the different imaging rounds.

3. To allow for accurate registration of the different masks for the same FOV, we corrected for

translation errors that emerge during imaging. For this we used the "Linear stack Alignment using SIFT" plugin under the registration option in ImageJ. This ensures that the centroids of the different cells in a FOV have roughly the same coordinates in the two masks. In our Python-based image registration code, we first determine the centroids of the cells in each mask separately. Next, we compare the euclidean distances between these centroids across the two masks. Since translational errors are corrected beforehand, the shortest distance between centroids in the two masks will correspond to the same cell. Therefore, this code helps identify the same cell across different imaging rounds and outputs its corresponding IDs from both masks. Thus at the end of registration we output a .csv file where for each FOV we have identified the cell ID of the same cells across the two masks.

4. In the final part of the image analysis, we wrote a python-based code to obtain the number of spots for different genes within each cells using its corresponding masks image and coordinates .csv file. Further we used the registration output to combine the spot counts across all rounds of FISH imaging for the same cells in the different FOVs.

The images across different time points were processed similarly to generate both the training and test files.

### S3 Analysis of spot count distributions across single cells.

#### 1. Negative Binomial fitting to the observed single-cell spot count data.

The RNA counts of single cells can be described by a negative binomial (NB) distribution which arises from the Telegraph model of bursty gene expression after certain approximations [4]. The underlying NB model can be parameterized by a burst size ($b$) and the burst frequency ($f$). The burst size $b$ refers to the number of RNA produced per activation event and the burst frequency $f$ signifies the number of such activation events per unit time. The probability of observing an smFISH count of $x_{ig}$ for gene $g$ in cell $i$, given a burst size $b_{ig}$ and burst frequency $f_{ig}$, is:

$$P(x_{ij}; b_{ij}, f_{ij}) = \frac{\Gamma(f_{ig} + x_{ij})}{\Gamma(x_{ig} + 1)\Gamma(f_{ig})} \left(\frac{1}{1 + b_{ig}}\right)^{f_{ig}} \left(\frac{b_{ig}}{1 + b_{ig}}\right)^{x_{ig}}, \tag{1}$$

where $\Gamma$ is the gamma function. The mean and variance of this NB distribution are given by $b_{ig}f_{ig}$ and $b_{ig}f_{ig} + b_{ig}^2 f_{ig}$ respectively.

We fit Equation (1) to spot count data for a given gene across cells for a particular time point using the `NonlinearModelFit` function in Mathematica. From the histogram of spot counts we extract the densities corresponding to the bin centers. The density *versus* bin center spot counts data is then fit using Equation 1 to obtain the best fit burst size and frequency. Similarly we fit the distribution over all the different training time points across the different clock genes. The NB distribution fits thus obtained for the different genes are shown in the different panels of Figure S3, and the burst size and frequency parameters are reported in Table S1 below. The main text Figure 1H shows plots of these parameters as functions of time.

| Genes → | *Bmal1* | | *Nr1d1* | | *Nr1d2* | | *Tef* | |
|---|---|---|---|---|---|---|---|---|
| Time points ↓ | $b$ | $f$ | $b$ | $f$ | $b$ | $f$ | $b$ | $f$ |
| 06 hrs | 6.81961 | 2.57057 | 42.392 | 1.08565 | 28.5877 | 1.20286 | 27.1325 | 1.3754 |
| 10 hrs | 10.6562 | 2.81861 | 33.6097 | 1.06534 | 8.40359 | 1.92026 | 10.9944 | 1.52946 |
| 14 hrs | 8.19197 | 3.48336 | 12.7258 | 1.86256 | 13.4216 | 1.4133 | 5.4954 | 1.69303 |
| 18 hrs | 16.6814 | 1.48567 | 83.7792 | 1.16196 | 15.8113 | 3.1386 | 20.2357 | 1.68867 |
| 22 hrs | 14.0021 | 1.09174 | 86.3773 | 1.11634 | 16.959 | 4.7936 | 19.9011 | 4.04528 |
| 26 hrs | 5.09502 | 1.56567 | 86.2862 | 1.01215 | 34.7216 | 2.83071 | 28.8422 | 2.81238 |
| 29 hrs | 5.11976 | 2.05427 | 93.0289 | 0.89279 | 21.2704 | 2.77934 | 33.1803 | 1.63852 |
| 34 hrs | 9.74369 | 2.16018 | 27.9645 | 1.03479 | 36.8863 | 1.17374 | 34.5223 | 1.0275 |
| 38 hrs | 10.6882 | 2.32803 | 22.4149 | 1.31782 | 27.8758 | 1.53469 | 17.0036 | 1.20196 |
| 42 hrs | 13.0018 | 1.88044 | 77.5818 | 1.01779 | 29.6518 | 2.55819 | 37.6699 | 1.09883 |
| 46 hrs | 14.0731 | 1.15072 | 47.9375 | 1.16443 | 13.1896 | 5.39904 | 22.248 | 2.60299 |
| 50 hrs | 8.73501 | 1.4187 | 31.9655 | 1.18606 | 19.5172 | 2.96948 | 31.1627 | 1.92063 |
| 54 hrs | 8.96564 | 1.66487 | 27.7884 | 1.27828 | 30.6301 | 1.88589 | 44.7296 | 1.28597 |

Table S1: Negative Binomial distribution parameters extracted from fits to SABER-FISH datasets for the different clock genes across time. $b$ refers to burst size and $f$ refers to the burst frequency.

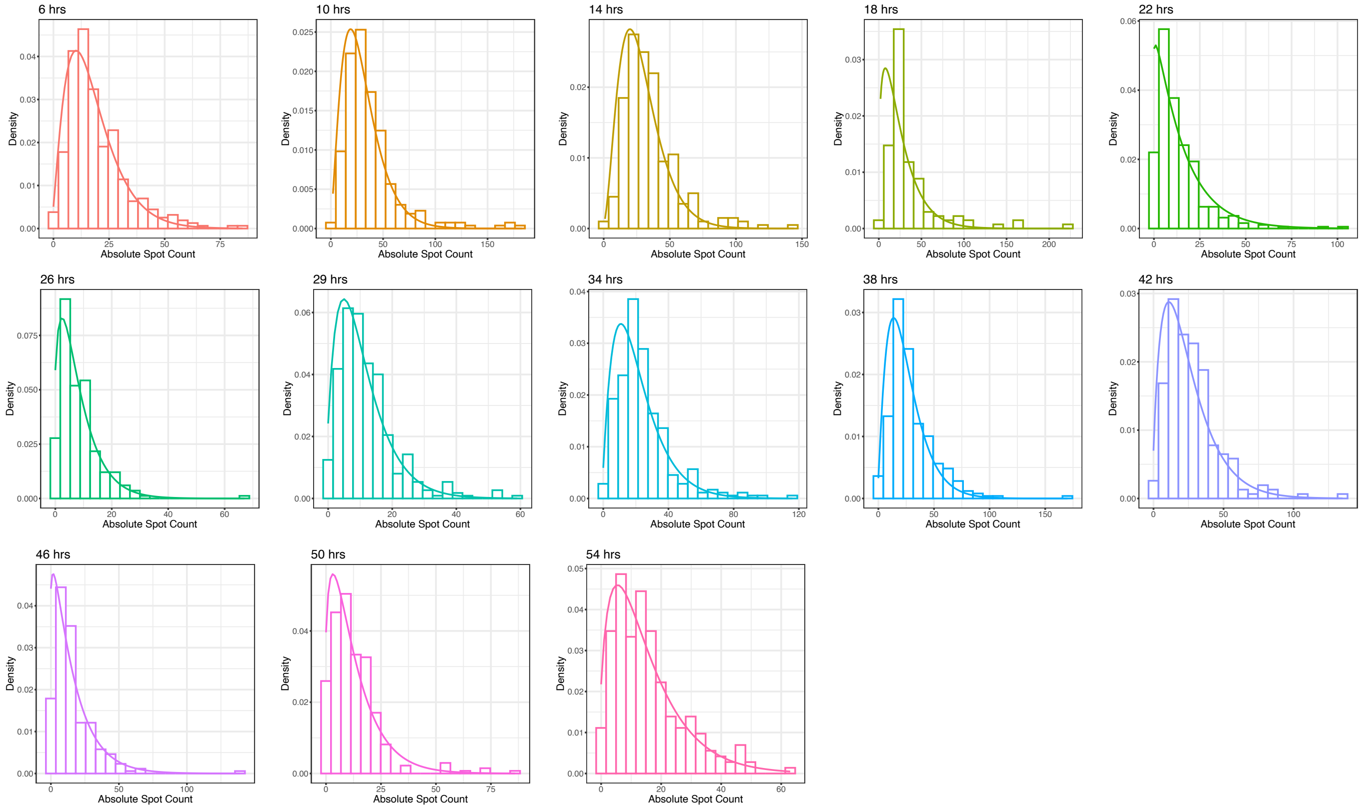

Figure S3: Negative binomial fits to the single-cell spot counts over the different training time points for *Bmal1* gene.

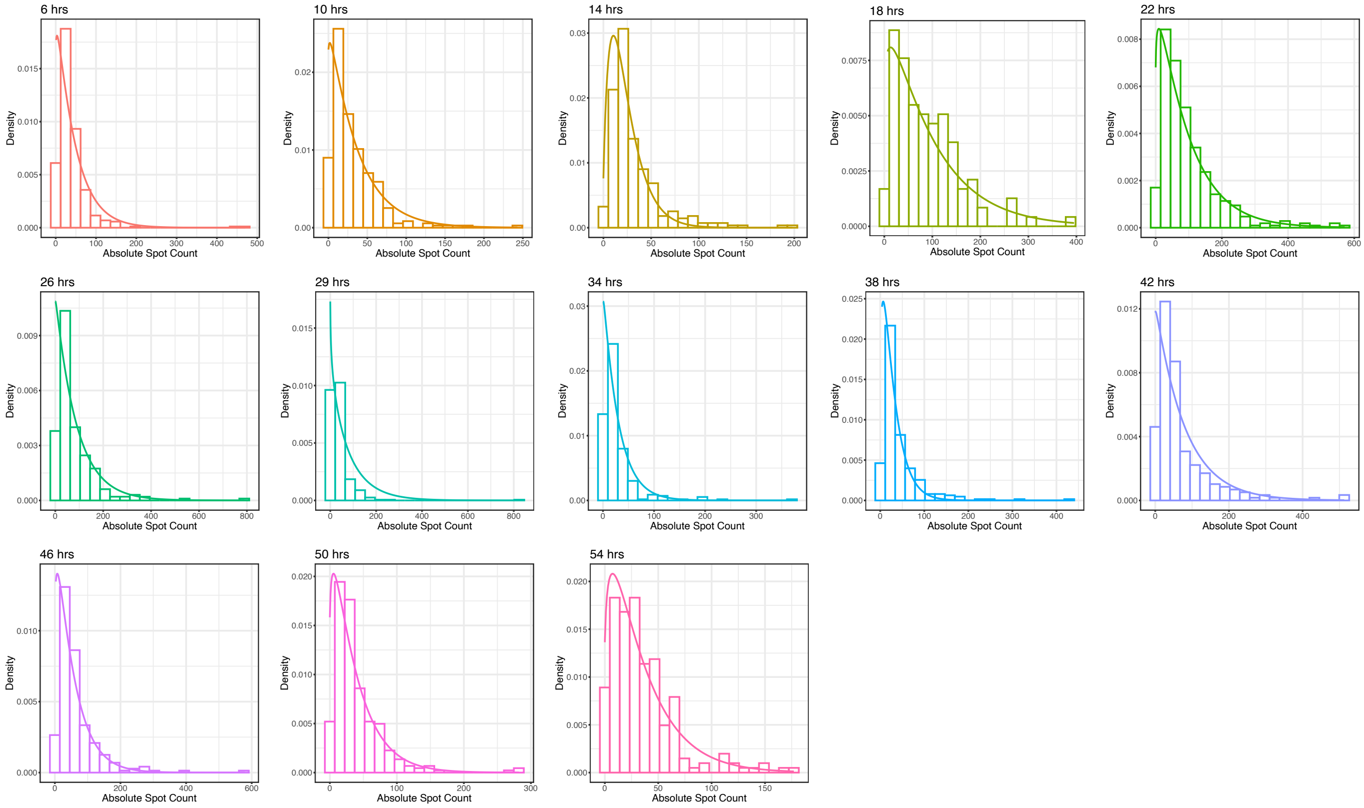

Figure S3: Negative binomial fits to the single-cell spot counts over the different training time points for *Nr1d1* gene.

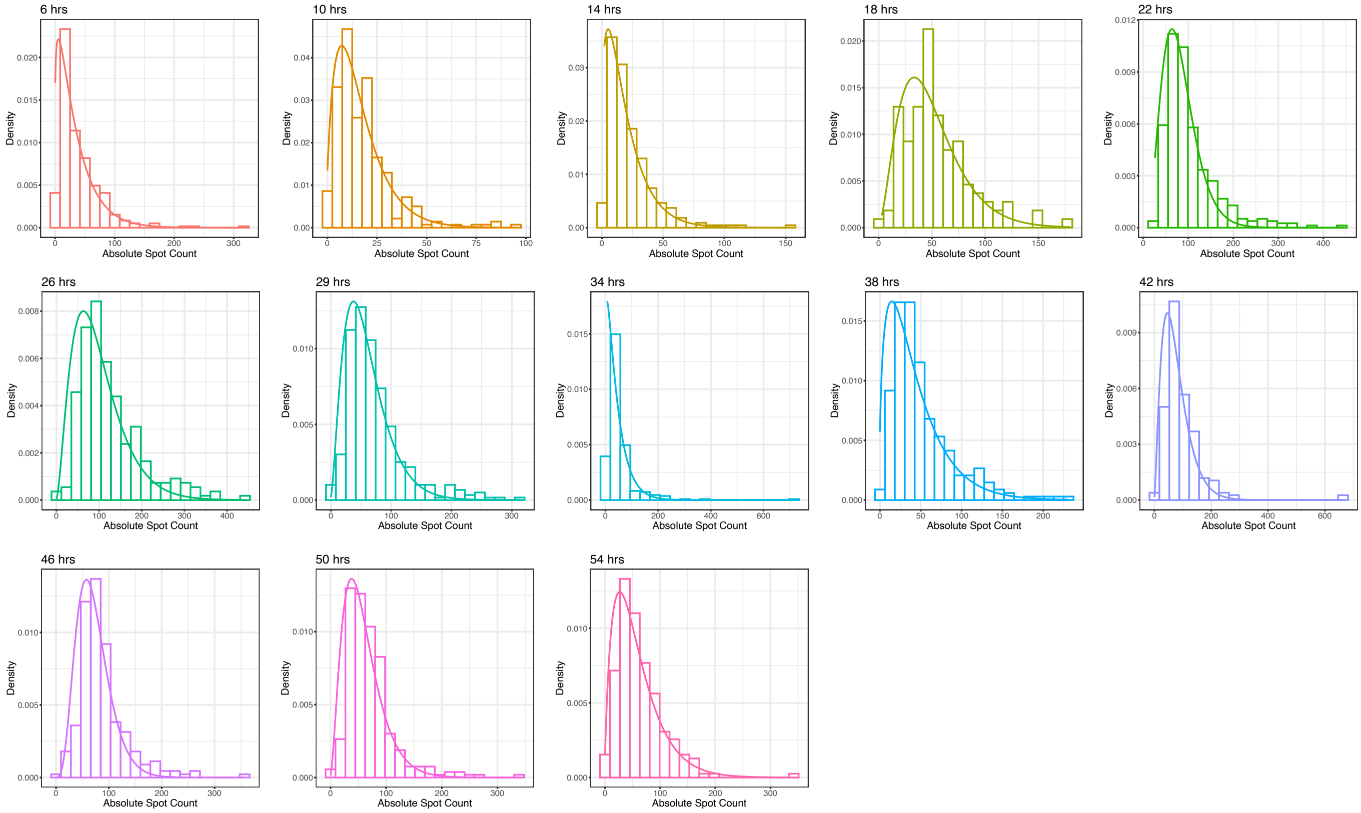

Figure S3: Negative binomial fits to the single-cell spot counts over the different training time points for *Nr1d2* gene.

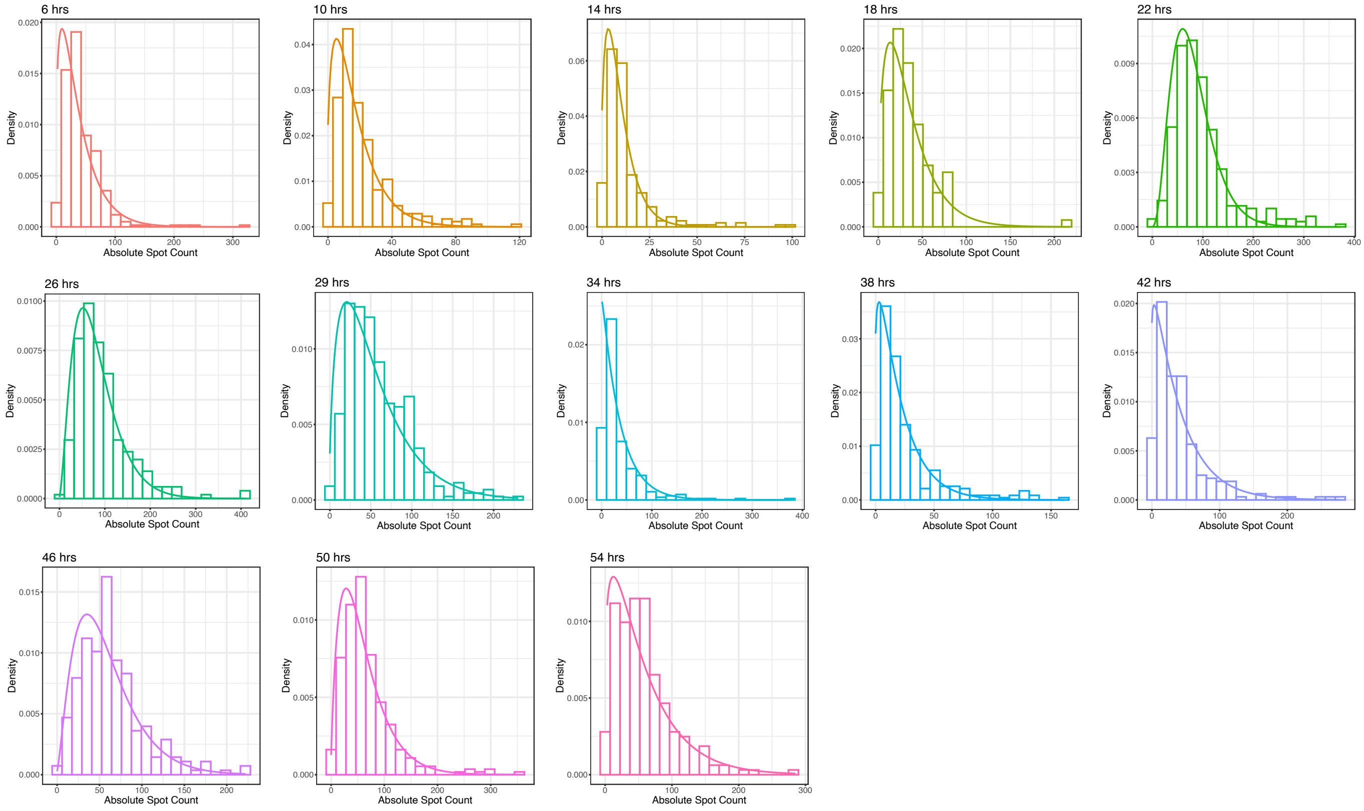

Figure S3: Negative binomial fits to the single-cell spot counts over the different training time points for *Tef* gene.

#### 2. Variance decomposition analysis of SABER-FISH datasets.

To check if we could detect variation in our datasets that could be ascribed to time, we performed variance decomposition following Bowsher and Swain [5]. In a biochemical system, if the fluctuations in the number of molecules of Z is determined by stochastic variables $Y_1$ and $Y_2$ and if $Y_1$ and $Y_2$ are conditionally independent then the total variation in $Z$ can be written as a sum of variations due to each stochastic control variable. Note that, $Y_1$ and $Y_2$ do not need to be the only sources of stochasticity in the system. The total variance in $Z$ is given by:

$$V[Z(t)] = V\{E[Z(t) \mid Y_1]\} + E\{V[E[Z(t) \mid (Y_1, Y_2)] \mid Y_1]\} + E\{V[Z(t) \mid (Y_1, Y_2)]\}, \qquad (2)$$

where the first and the second terms represent the 'explained' variation in $Z$ due to $Y_1$ and $Y_2$ respectively, whereas the third term quantifies the 'unexplained' variation due to sources other than $Y_1$ and $Y_2$. We applied Equation (2) to quantify the variation in RNA counts due to the two random variables – Cell Area, and Time. In our analysis, for all 4 genes, Time $\in \{6, 10, 14, ...54\}$ and Area was normalised and binned before conditioning with bin size of $10^{-4}$. Since there is no unique way of assigning area and time to $Y_1$ and $Y_2$, we calculated the variance explained by each random variable by taking the average between the following two cases: first considering area as $Y_1$ and time as $Y_2$ and then the other way around. For example, for a given gene we first conditioned the RNA count on area and calculated the variance of the mean (across all the area bins), $V\{E[\text{RNA count} \mid \text{area}]\}$. Then we conditioned the expectation of RNA count given time and area upon area, and calculated the mean of the variance given by $E\{V[E[\text{RNA count} \mid (\text{area, time})] \mid \text{area}]\}$. The average of these two terms, $V\{E[\text{RNA count} \mid \text{area}]\}$ and $E\{V[E[\text{RNA count} \mid (\text{area, time})] \mid \text{area}]\}$ therefore represents the variation in the data due to area. The contribution from time was calculated similarly. The fraction of total variance explained by the different sources as computed by this approach is listed in Table S2. The total variance calculated was slightly lower than the total variance calculated from the raw data. This limitation is likely due to the low sample numbers we had after conditioning on area or time.

| Gene | Area | Time | Other sources |
|---|---|---|---|
| *Bmal1* | 0.76 | 0.22 | 0.02 |
| *Tef* | 0.72 | 0.26 | 0.02 |
| *Nr1d1* | 0.80 | 0.18 | 0.02 |
| *Nr1d2* | 0.77 | 0.22 | 0.01 |

Table S2: Fraction of total variance explained by Cell Area, Time and other unknown sources

#### 3. ODeGP-based oscillation detection for the different clock genes

ODeGP is a classification algorithm that combines Gaussian Process regression and Bayesian inference to sensitively detect the presence of oscillations in noisy, non-stationary time-course datasets [6]. In order to analyze the oscillations present in our training dataset, we performed bootstrapping over the cells included in our data over the different time points. For a specific gene we randomly selected a group of 100 cells from a specific time point to obtain the mean gene expression. We repeated this 50 times for each of the training time points in order to generate the means and errors of the expression data. Similarly we obtain the mean and errors for the spot counts for all the clock genes considered.

Each of the genes were separately analyzed using the `oscOrNot` function in the `OdeGP` package in R with the arguments : nonstat = TRUE, threshold = 14, detrend = FALSE, plotting = TRUE, Q = 1 to detect the presence of oscillations.

#### 4. Comparison of burst size and frequency over the different training time points.

To identify which parameters of transcriptional bursting gave rise to the observed oscillations of the clock genes, we compared the burst size and frequency of the different genes across the training time points as shown in main text Figure 1H. We plotted the burst size and frequency extracted in Section 3.1 as a function of time post Dex-resetting to visualize the trends in the parameter estimates.

### S4 Bulk gene expression measurements using qPCR.

#### 1. Experimental protocol and analysis.

To obtain the bulk gene expression measurements for the clock genes – *Bmal1* and *Nr1d2*, we performed a Quantitative real-time PCR (qPCR) assay. Similar to the synchronisation protocol followed for SABER-FISH experiment, we seeded cells at two different seeding density of 15000 cells/well and 30,000 cells/well in 16 different wells across multiple 6-well plates. Time points – T3, T6, T9, T12, T15, T18, T21 and T24, were collected at the same time while time points T27, T30, T33, T36, T39, T42, T45 and T48 were collected at a different time. After allowing the cells to attach for 12-15 h, we began Dex-synchronising the cells in different wells using the same protocol as mentioned in section S1 every 3 h. In order to eliminate the initial transients of Dex treatment, we refrained from collecting data in the initial 3 h post synchronisation hence the time points ranged between 3 to 48 h. At the end of the 48 h we extracted the cells using Trizol and proceeded to extract the total cellular RNA using Purelink RNA extraction kit (Invitrogen, cat. no. 12183018A). After RNA extraction, we performed cDNA conversion of cellular RNA using SuperScript III First-Strand Synthesis System (Invitrogen, cat. no. 18080051) with oligo-dT as the primer. Finally, qRT-PCR was performed in $10\mu$L final reaction volume, with three replicates per sample, by using iTaq Universal Sybr-Green supermix (Biorad, cat. no. 1725121). We added 15 ng of cDNA per well and used a final primer concentration of 400 nM for qRT-PCR. We also kept three non-template controls (NTC) for each gene as our negative controls.

After obtaining the Ct values for the different genes in the different time points, we performed $\Delta\Delta$Ct analysis to obtain the fold change of the clock genes with respect to *Gapdh*. For the $\Delta\Delta$Ct analysis we consider Ct values of the *Gapdh* gene as the endogenous control and the 3 h sample as our reference control. Post normalization with respect to the endogenous and reference control, we express the Ct values as fold change.

The fold change with respect to *Gapdh* of the different clock genes as a function of time post Dex-resetting was subjected to OdeGP-based oscillation classification. Each of the clock genes' data were separately analyzed using `oscOrNot` function in the `OdeGP` package in R with the arguments :

nonstat = TRUE, threshold = 14, detrend = TRUE, plotting = TRUE, Q = 1 to detect the presence of oscillations.

##### 2. Primer sequences used in the qPCR experiments.

The primer sequences used for performing the qRT-PCR of the clock genes – *Bmal1* and *Nr1d2* and the consitutively expressed gene *Gapdh* are mentioned in Table S3.

| Primer Label | Sequence |
|---|---|
| *Gapdh* Forward Primer | AACTTTGGCATTGTGGAAGG |
| *Gapdh* Reverse Primer | GGATGCAGGGATGATGTTCT |
| *Bmal1* Forward Primer | GGGCTGGACGAAGACAATGA |
| *Bmal1* Reverse Primer | CGCCCGATTGCAACGA |
| *Nr1d2* Forward Primer | CCCAAGAACGCTGATATCTCTAG |
| *Nr1d2* Reverse Primer | ACACAGTAGAACCATGCCAC |

Table S3: qPCR primers

#### S5 Simulations to explore the role of averaging on noise reduction.

We generated pseudo-time trajectories using the procedure as mentioned in main text for the different clock genes included in our training dataset. We observed that when the expression was averaged over a group of only 10 cells, we observed clean oscillations as shown in Figure 2A and Figure S4. In order to gain intuition how averaging over cells might lead to reduction in noise and emergence of clean oscillations we performed the following simulations:

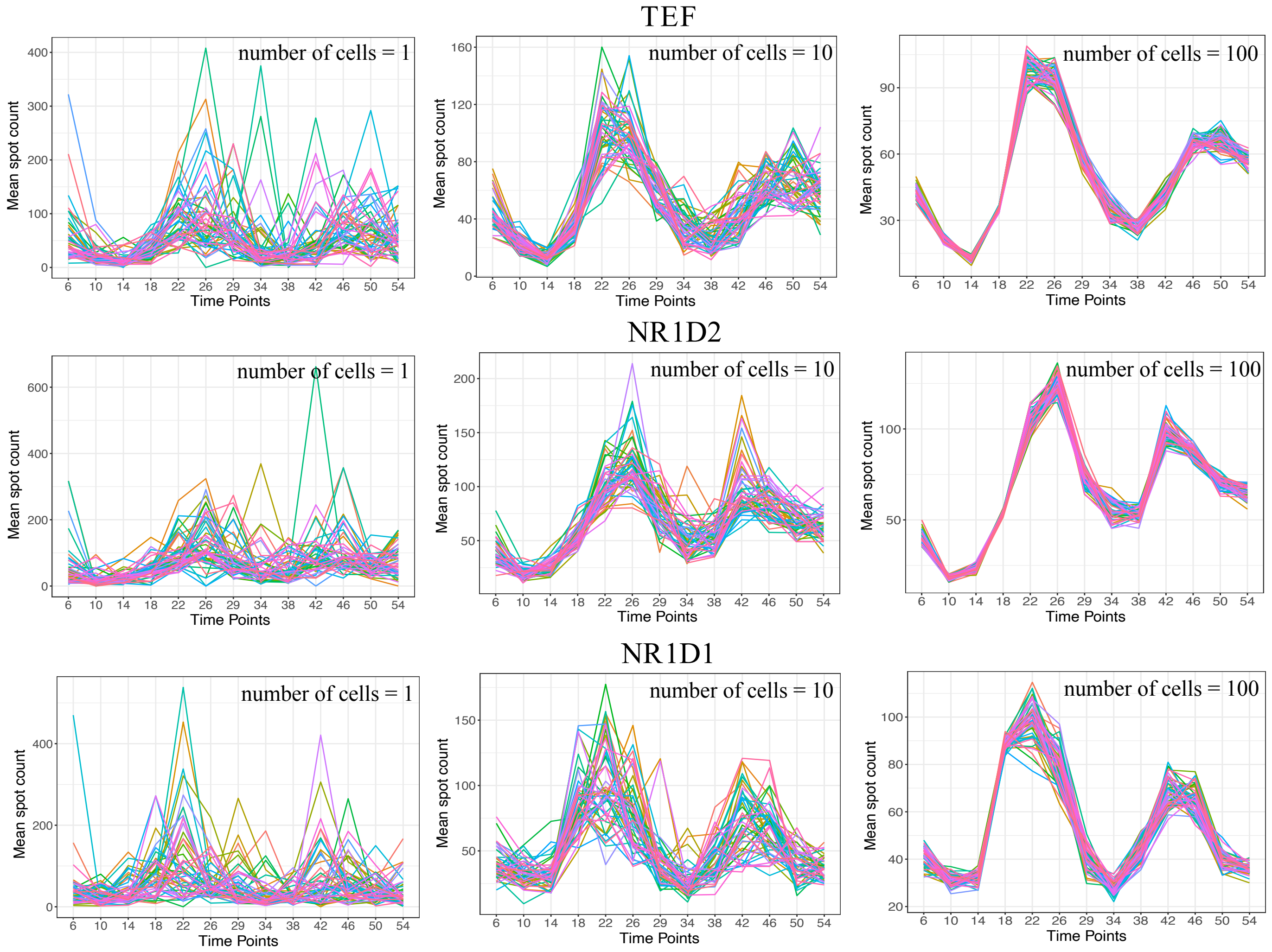

Figure S4: Pseudo-time trajectories for the clock genes included in the NIH3T3 training dataset. Trajectories for *Bmal1* are shown in the main text Figure 2.

### 1. Simulating noisy sine functions.

In order to generate a single noisy trajectory, we utilized the `sin` function in R to obtain the values of the sine curve with a time period of 24 h at every 1 h interval over 48 h and added Gaussian noise with mean and standard deviation for the different cases as mentioned in Table S4. We simulated multiple (N = 300) such noisy sinusoidal curves for each of the following noise levels, representing individual cells. We then averaged over different cell numbers for each of the cases to compare the population level oscillation with the individual cellular oscillation.

| Noise level | Mean | Standard Deviation |
|---|---|---|
| Low | 0 | 1.5 |
| Medium | 0 | 5 |
| High | 0 | 10 |

Table S4: Parameters of the Gaussian noise added to the sinusoidal trajectories

#### 2. Generating synthetic spot count data from NB distribution fits.

To obtain a more realistic representation of the underlying noise in clock gene expression, we generated synthetic spot count data of the different clock genes – *Bmal1*, *Nr1d1*, *Nr1d2* and *Tef* at the different training time points by sampling from Negative Binomial distributions with parameter values obtained from the fitting performed as described in Section S3. As mentioned in Section S3, we obtain the burst size (b) and the burst frequency (f) that parameterizes the underlying Negative Binomial at a particular time point for a specific gene. The mean and variance of this underlying NB distribution depends on the burst size and the burst frequency as follows:

$$\mu = bf \tag{3}$$

$$\sigma^2 = bf + b^2 f \tag{4}$$

In an alternative parameterization, the Negative Binomial distribution is specified by the number of successes (denoted by $r$) and the probability of success in each trial (denoted by $p$). In this parameterization, the distribution is given as follows:

$$P(k; r, p) = \binom{k+r-1}{k} (p)^r (1-p)^k \tag{5}$$

where $k$ is the number of failures or in other words $k+r$ trials are required to accumulate $r$ successes. In this form of the NB distribution, the mean and variance are given by:

$$\mu = \frac{r(1-p)}{p} \tag{6}$$

$$\sigma^2 = \frac{r(1-p)}{p^2} \tag{7}$$

This parameterization has been adopted in the `stats` package of R which we use to generate the synthetic spot counts by sampling. Hence we derive the parameters $r$ and $p$ in terms of the best fit burst size and frequency ($b$ and $f$) to generate the synthetic spot counts.

$$r = f \tag{8}$$
$$p = \frac{1}{1+b} \tag{9}$$

At each time point for the *Bmal1* gene we sampled 200 random numbers from a NB distribution using the `rnbinom` function with the parameter values : n = 200, size = r, prob = p. After generating the single-cell spot counts for the different time points, we generated pseudo-time trajectories as described in main text. This procedure was carried out for all the genes.

### S6 PRECISE: Predicting Circadian phase from Stochastic gene Expression

#### 1. Bootstrapping to calculate mean expression and errors of the mean.

In order to generate the training data from single-cell spot counts, we bootstrapped the data from each time point to generate mean expression levels and errors of the mean. For a specific gene at a particular time point, we randomly sampled 100 cells and obtained the mean expression value. We repeated this sampling procedure (with replacement) 50 times to generate 50 mean expression values. This provided the mean of means and error of means. The entire procedure was repeated for each gene and each time point to generate the bootstrapped data for all 4 genes across all the 13 different time points. Note that unlike the distribution of spot counts which is Negative Binomial, the distribution of mean spot counts is very close to being normally distributed (*Bmal1* distributions of the mean are shown in main text Figure 5F). As discussed in later sections, the Gaussian Process based algorithm ODeGP is always used on this training data comprising the means and errors of the means.

### 2. Wavelet based true phase assignment of training and test samples.

We next converted the time labels of the training and test samples to circular phase values. As described in section S1 and S2, we collected data over different times post Dexamethasone treatment for both training and test samples, corresponding to distinct circadian phases. To assign phases to each of the samples (training and test) collected at different time points, we determined the phase of each sample using pyBOAT, an algorithm that uses wavelet transform to assign phases to non-stationary (the period and amplitude vary over time) oscillatory data.

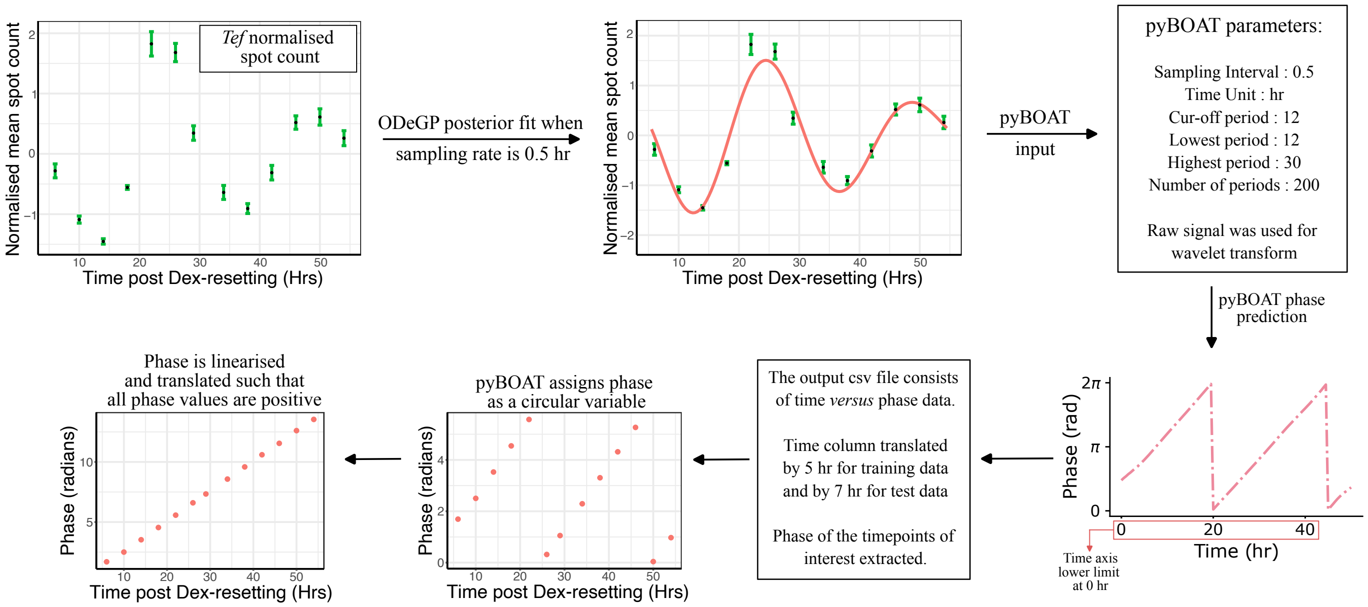

Figure S5: An overview of pyBOAT based true phase assignment of training and test samples : Example has been shown when *Tef* gene is used as input to assign the phase for both training and test samples.

The circadian phases of the training samples ($\phi_i$, where $i = 1, 2, ..., 13$) can be estimated from the Dex-synchronized time-series of normalized spot counts of any oscillatory gene included in the data using pyBOAT. The bootstrapped spots counts are normalized discussed in Section S6.3 and then these normalized mean spot counts are provided as input to pyBOAT for obtaining the circular phase values for the samples. However pyBOAT takes as input data sampled at regular intervals while our datasets, especially the test time points, comprised irregularly spaced time points. To overcome this problem, we utilised our previously developed algorithm ODeGP to determine the posterior mean fit to the data points and used that as input for pyBOAT. ODeGP is a classification algorithm that determines whether input data is oscillatory or not. In parallel, it also provides a mean fit to the data points. The ODeGP fit becomes progressively worse as we try to estimate it for time points that are further away from the actual observed time points, hence for training data we estimated the mean fit in the range of 5-55 h at every 0.5 h interval, and for the test data the mean fit was determined in the range 7-33 h with 0.5 h interval between consecutive time points. As mentioned previously, this fit is used as input to pyBOAT along with the sampling interval (0.5 h) and parameter values as mentioned in Figure S5 to determine the phases. However, since the absolute time values in pyBOAT are considered from 0 h, before extracting the true phase we translate the time column of the output file by 5 h for training data and 7 h for test respectively and then obtain phases at our time points of interest. Since phase is a circular variable and pyBOAT assigns it between 0 and $2\pi$, we made it linear by subtracting or adding $2\pi$ as necessary and also made sure that at the end all the phase values were positive. This step was necessary for application of ODeGP to the data as mentioned later (Figure S5).

Finally, an important consideration was that the wavelet-based phase inference software pyBOAT suffers from edge effects – for the end samples beyond which observed data points are absent on either side, the phase cannot be assigned accurately [7]. To account for this issue, during validation of PRECISE's performance, the time points on the ends, that is 8 and 32 h, were excluded from all analysis.

After this step, our final training data (used for all downstream purposes) comprised the mean spot counts of the different genes along with the errors of the means as functions of circadian phase. The circadian phases of the 13 different time points are denoted by $\phi_i$, where $i = 1, 2, 3....13$ through the rest of this text. Along with the training data, the above steps also provided the estimates of true phases of the different test samples ($\phi_{true}$). The performance of the different phase prediction algorithms were checked in terms of deviations of the predicted phase from these true phase estimates ($\phi_{true}$) of the different test samples.

#### 3. Normalization of data to allow estimation of circadian phase across experiments and cell types.

Systematic differences across experiments conducted at different times or in gene expression of different cell-types requires normalization of datasets. Therefore to allow prediction of circadian phase across cell types and data collected across different experiments, we performed the following data normalization procedures on both training and test datasets (all vectors in the following sections are denoted by boldface letters):

##### 3. .1 Normalization of training data.

For our training data we have $i$ different circadian phases ($i = 1, 2, ...13$) and $j$ different genes ($j = 1, 2, 3, 4$). Upon bootstrapping, for a single gene $j$ we generated 50 mean samples (each is a mean expression over 100 cells) for every phase $i$. Let $\mu_{ij}$ and $\sigma_{ij}$ be the mean and standard deviation over the 50 samples belonging to the $i^{th}$ circadian phase for the $j^{th}$ gene. For the $j^{th}$ gene, consider $\mathbf{g_j}$ to be a 13×1 vector that includes the mean $\mu_{ij}$ over the $i$ different phases. For this vector $\mathbf{g_j}$, we calculate the mean and standard deviation over the $i$ different phases as follows :

$$\mu_j = \frac{\sum_{i=1}^{13} \mu_{ij}}{13} \tag{10}$$

$$\sigma_j = \sqrt{\frac{\sum_{i=1}^{13} (\mu_{ij} - \mu_j)^2}{13}} \tag{11}$$

For the $i^{th}$ phase, we normalize the mean ($\mu_{ij}$) and standard deviation ($\sigma_{ij}$) as follows :

$$\mu_{ij}^{'} = \frac{\mu_{ij} - \mu_j}{\sigma_j} \tag{12}$$

$$\sigma_{ij}^{'} = \frac{\sigma_{ij}}{\sigma_j} \tag{13}$$

We separately perform the above normalization for the different clock genes. This ensures that mean and standard deviation of the spot counts for each of the clock genes over the different circadian phases is 0 and 1 respectively.

##### 3. .2 Normalization of test data.

Similar to the training data we generate bootstrapped data for the test samples as well. Thus for each of the test time points, we consider 50 samples, where each sample is characterised by spot count averaged over randomly chosen fixed number of cells. Similar to normalization performed for the training data, we obtain the values of $\mu_j$ and $\sigma_j$ that represent the mean and standard deviation of mean spot count data over the test time points. We use the values of $\mu_j$ and $\sigma_j$ to individually normalize each of the 50 samples across each of the different test time points according to equation 12. While we needed at least 3 time point samples in our test data for this normalization, the actual phase prediction were performed for snapshots where each test time point's inference was done separately when its normalized clock genes counts were provided as input to PRECISE.

#### 4. Training PRECISE to learn the posterior mean and variance.

The training data input is the normalized mean RNA counts $\mu_{ij}^{'}$ of different genes as a function of circadian phase $\phi_i$ assigned using pyBOAT. Training involved subjecting normalized mean spot count *versus* phase data for each gene to ODeGP based classification as oscillatory or non-oscillatory. As mentioned, ODeGP uses non-stationary kernel to describe oscillatory functions, hence if a gene is classified as oscillatory PRECISE records the corresponding optimised hyper-parameters of the non-stationary kernel and performs this separately for each gene. The OdeGP classification for the different genes were performed using the `oscOrNot` function with the arguments : nonstat = TRUE, threshold = 14, detrend = FALSE, plotting = TRUE, Q = 1, to detect the presence of oscillations. For the $j^{th}$ gene we denote the vector of optimised hyper-parameters by $\boldsymbol{\theta_j}$

#### 5. Test sample phase prediction.

For the $j^{th}$ gene, consider a 13×1 normalized mean vector $\mathbf{g_j^{'}}$ comprising normalized mean values $\mu_{ij}^{'}$. The data of this normalized count vector $\mathbf{g_j^{'}}$ as a function of the training phases $\phi_i$ is classified using ODeGP and the optimised hyper-parameters (denoted by $\boldsymbol{\theta_j}$) of the non-stationary kernel are recorded. If the test sample has a normalized count of $g_{j*}$ for the $j^{th}$ gene, then the joint distribution of the observed function values $\mathbf{g_j^{'}}$ at the training phases $\phi_i$ and the function value $g_{j*}$ at the test phase $\phi_*$ (unknown at the moment) under the assumed prior is given as:

$$\begin{bmatrix} \mathbf{g_j^{'}} \\ g_{j*} \end{bmatrix} \sim \mathcal{N}\left( \begin{bmatrix} \mathbf{m_j} \\ m_{j*} \end{bmatrix}, \begin{bmatrix} K_{\phi_i\phi_i} + \sigma_{n^2} I & K_{\phi_i\phi_*} \\ K_{\phi_*\phi_i} & K_{\phi_*\phi_*} \end{bmatrix} \right)$$

where,

$\mathbf{m_j}$ = mean vector for the training gene expression values at the different training phases

$m_{j*}$ = mean vector for the test gene expression value at the test phase

$K$ = covariance matrix whose entries are calculated using the non-stationary kernel

Then the posterior probability distribution of $g_{j*}$, given the training gene expression values $\mathbf{g_j^{'}}$, training phases $\phi_i$ and the non-stationary kernel hyper-parameters $\boldsymbol{\theta_j}$ is given by

$$P(g_{j*}|\phi_*, \phi_i, \mathbf{g_j^{'}}, \boldsymbol{\theta_j}) = \mathcal{N}(\mu_{j*}, \sigma_{j*}^2) \tag{14}$$

where

$$\mu_{j*} = m_{j*} + K_{\phi_*\phi_i}[K_{\phi_i\phi_i} + \sigma_{n^2} I]^{-1}(\mathbf{g_j^{'}} - \mathbf{m_j}) \tag{15}$$

$$\sigma_{j*}^2 = K_{\phi_*\phi_*} - K_{\phi_*\phi_i}[K_{\phi_i\phi_i} + \sigma_{n^2} I]^{-1} K_{\phi_i\phi_*} \tag{16}$$

We can similarly define the posterior probability distribution for the expression values of all the $j$ genes in the test sample. Then the total data likelihood ($L$) and the Total *log* likelihood ($TLL$) can be written as

$$L = \prod_{j=1}^{4} P(g_{j*}|\phi_*, \phi_i, \mathbf{g_j^{'}}, \boldsymbol{\theta_j}), \tag{17}$$

$$TLL = \log(L) = \sum_{j=1}^{4} \log(P(g_{j*}|\phi_*, \phi_i, \mathbf{g_j^{'}}, \boldsymbol{\theta_j})). \tag{18}$$

In the above expression we know all the values except the test phase $\phi_*$. We constructed the total likelihood $L$ *versus* phase graph to determine the predicted phases of the test samples. We detect the phases corresponding to the local maxima in $L$ followed by taking the circular mean over these phases to obtain the average phase prediction $\phi_{pred}$. Since our training data spanned over 48 hr corresponding to two circadian cycles, we expect multiple peaks during phase prediction and hence an average phase estimate is more accurate. We use the `findpeaks` function in the ggpmisc R package with arguments : span = 3, ignore_threshold = 0.3 to find the positions of local maxima. For computing the circular mean, we use the `mean.circular` function in the circular package in R [8].

Thus, upon optimisation the algorithm predicts the phase of the test sample characterised by a specific set of gene expression values.

#### 6. Calculation of deviation between PRECISE-based phase prediction and true phases.

Since our test samples were also derived from synchronised cells, we were able to assign the true phase of these test datasets using pyBOAT ($\phi_{true}$) as mentioned in Section 6.2. Thus we compared the PRECISE inferred phase ($\phi_{pred}$) and the true phase to quantify the accuracy in prediction. The deviation in prediction from the true phase for each test sample, $\Delta$, was calculated as follows:

$$\delta = |\phi_{pred} - \phi_{true}| \tag{19}$$

$$\Delta(Hrs) = \begin{cases} \delta * (24/2\pi), & \text{if } \delta \leq \pi \\ |2\pi - \delta| * (24/2\pi), & \text{if } \delta > \pi \end{cases}$$

Using the above expression we evaluate the absolute deviation in phase estimate whose value lies between 0 - 12 h. To obtain the average deviation from true phase for samples where multiple peaks are detected in the likelihood $L$ *versus* phase graph, we calculated the deviation from true phase separately for the phases corresponding to the different peaks and took a mean of the deviations thus obtained.

In order to determine the limits imposed by biological noise with regards to phase inference, we first began by estimating the deviations obtained for the NIH3T3 test samples using different combinations of two clock genes, when different clock genes were used to assign the true circadian phase (Figure S6).

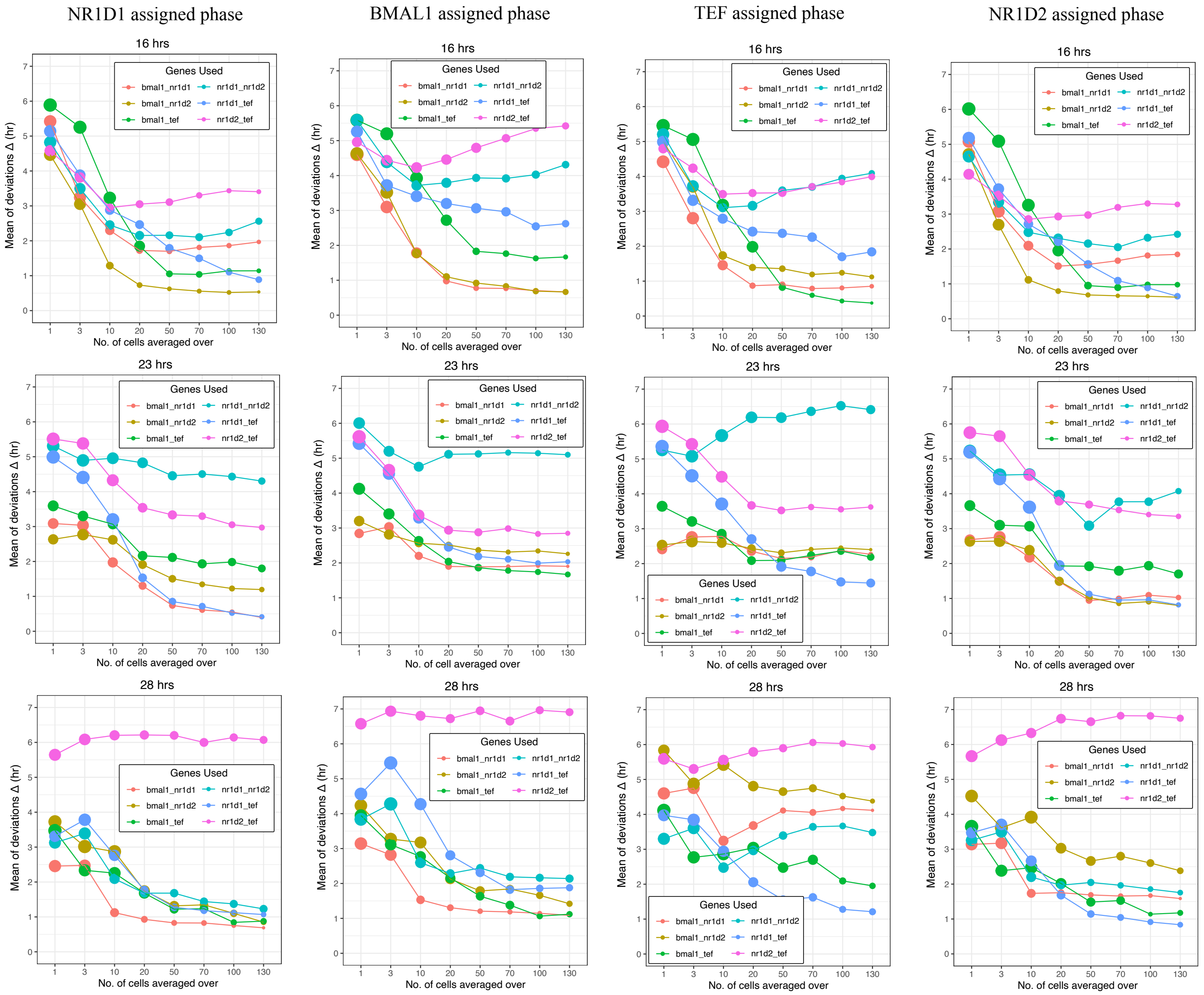

Figure S6: PRECISE-based phase predictions using 2-gene combinations, when different genes were used to assign the true phases. For each of different genes considered to assign true phase, we also compared the deviations across different possible combinations of two clock genes across the three different test time points included in NIH3T3 data. In most cases, phase inference using 2-gene combinations produced large deviations, except for the 28 hour sample with *Nr1d1* assigned true phase.

Since the error associated with most of the two gene combinations were high, we next explored the performance of three clock genes when used for phase inference. We visualized the deviations both in the form of boxplots and also empirical cumulative distribution function (eCDFs). As shown in main text Figure 4A and Figure S7, while the deviations span the entire 12 h when phase is estimated for gene counts of true single cells, upon averaging the gene expression over small cellular groups, we see a decrease in deviation to less than 1 h which saturates after a specific size of the cellular group, further quantified by the minimum performance improvement for higher sample sizes in the eCDFs of the deviations.

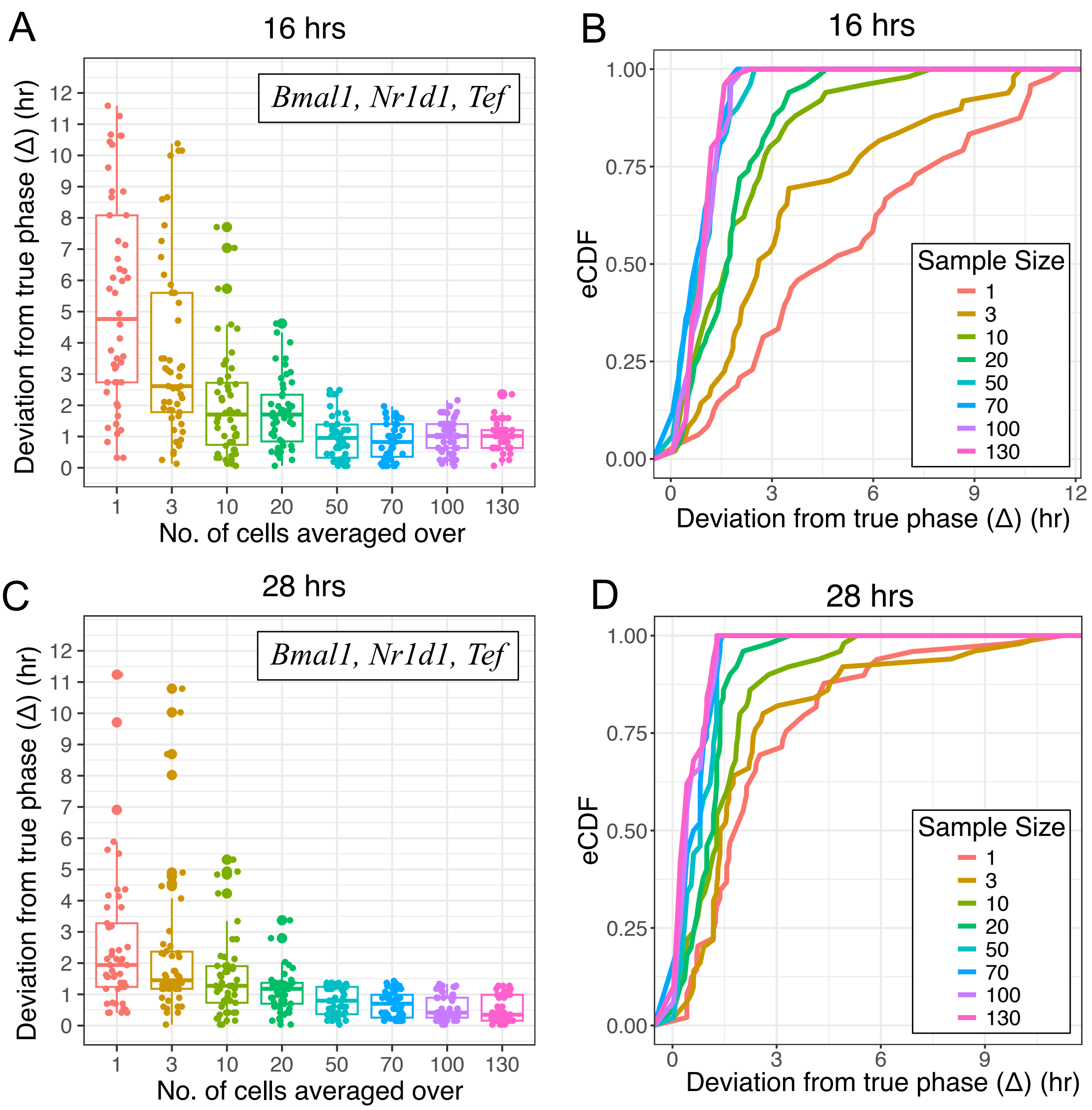

Figure S7: PRECISE-based phase predictions using the 3 genes *Bmal1*, *Nr1d1* and *Tef* on NIH3T3 cells. Here *Nr1d1* was used to assign the true phases. The 16 and 28 hour test samples are shown here; the 23 hour sample is shown in the main text Figure 4A.

We also explored the performance of the other combinations of three clock genes against using all four genes for prediction when different genes were used to assign true phases. As shown in Figure S8 irrespective of the gene used to assign true phase, a three gene combination performed as well as using all the four genes across the three different time points in NIH3T3 data.

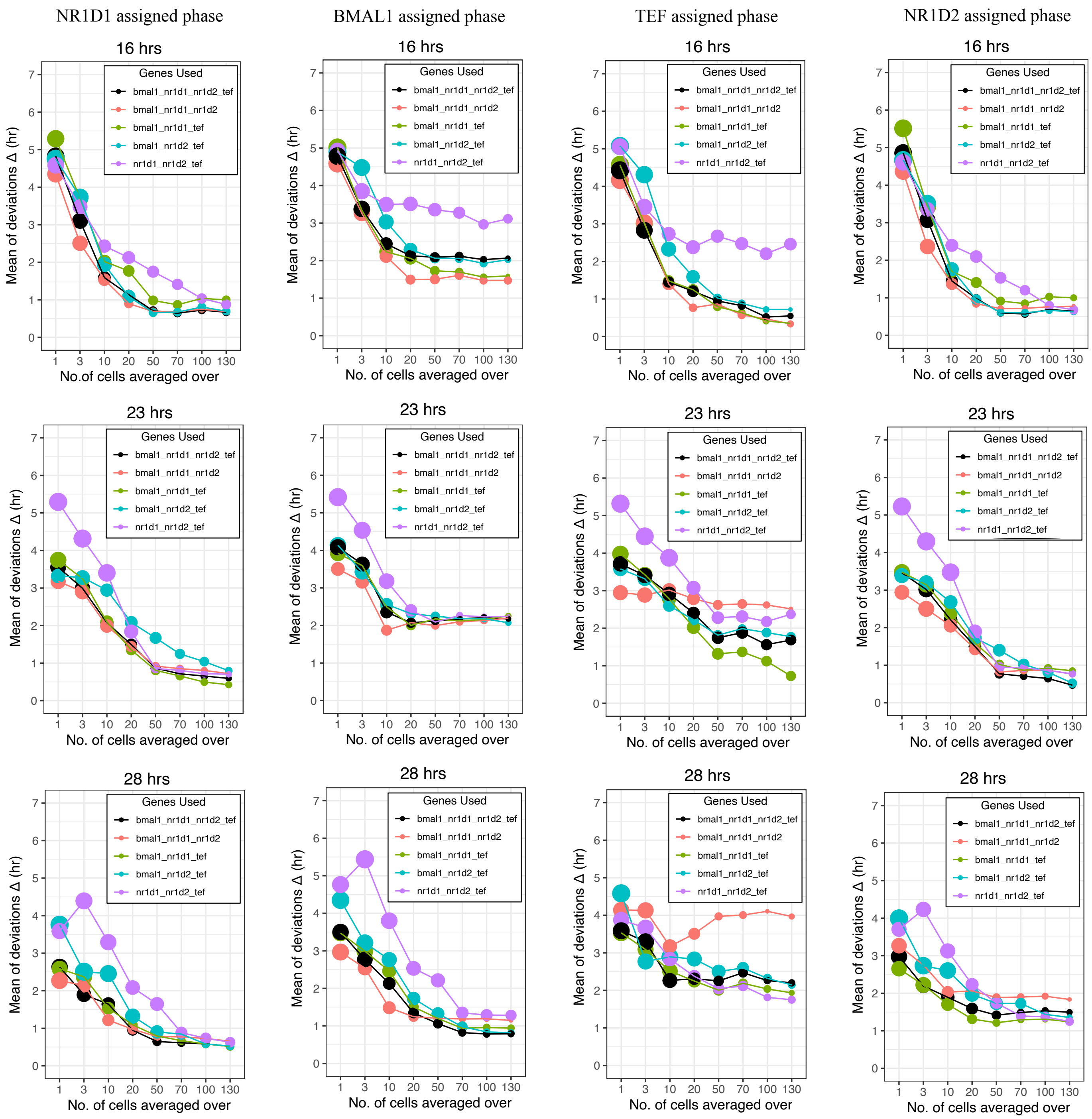

Figure S8: PRECISE-based phase predictions when different genes were used to assign the true phases. For each of different genes considered to assign true phase, we compared the deviations across different possible combinations of 3 clock genes and also all 4 genes across the three different test time points included in the NIH3T3 data.

### S7 Circadian phase estimation using Singular Value Decomposition.

We compared the prediction of our supervised learning algorithm PRECISE with Singular Value Decomposition (SVD), which is an unsupervised learning algorithm to order samples according to phase. SVD-based circular phase inference works on the principle that oscillatory gene expression will result in a high dimensional ellipse (Main text Figure 1A). The major axis of this ellipse represents the direction of maximum variance in the data, therefore identifying this direction is equivalent to finding the first principal component (first eigenvector of the covariance matrix). The minor axis will be the second principal component. Hence with SVD based dimensionality reduction to two dimensions and projection of samples onto this 2D eigenvector space, the elliptical structure will be preserved along with relative ordering of the samples. This allows quantitative retrieval of phase information of each sample.

As described in the main text we generated a 3×250 matrix whose entries were expression values of 3 clock genes (*Bmal1*, *Nr1d1* and *Tef*) in 50 randomly chosen single cells from each of the 5 different time points. We normalized the entries of the matrix as described in Section S6 such that the mean and variance for the expression of the clock genes over the different time points was 0 and 1 respectively. This matrix was subjected to SVD and then the samples (denoted by the columns of the 3×250 matrix) were projected onto the first 2 eigengene space. A similar procedure was repeated to obtain the projection of the 3×250 data matrices when the entries were the gene expression averaged over randomly selected groups of cells. We averaged over groups containing 3, 10, 20, 50, 70, 100 and 130 cells respectively as explained in the main text.

#### 1. Calculation of deviation between SVD-based phase prediction and true phases.

In order to obtain circadian phases of the samples from the SVD projections, we calculated the $tan^{-1}$ of the ratio of the coordinates in the eigengene space. For this we utilised the `atan2` function included in base R package that determines the phase in terms of radians. To compare the performance of

PRECISE-based phase inference with respect to SVD, we compared the SVD-based inferred phase with the true phase estimates assigned using pyBOAT (Section S6). As mentioned in Section S6, the true phase assignment was performed with respect to a particular clock gene, since the peak of any gene can be arbitrarily defined as phase 0 (or $2\pi$). However, this was not directly comparable to the phase inferred using SVD and thus we needed to make the two estimates comparable.

As shown in Figure S9, when higher number of cells are selected to obtain the average gene expression, samples belonging to the different test time points are clustered closely together and hence the phase assigned for these samples can be considered to be more faithful representation of the true circadian phase. Thus from the SVD projection of the matrix comprising gene expression averaged over 130 cells, we obtain the mean of the inferred phases across the 50 samples per time point separately for each test time point. Thus, we obtain 5 mean SVD inferred phases for the 5 different test time points included in our NIH3T3 data. We set the phase of the 8 h test sample to be zero in both SVD' prediction as well as pyBOAT's assignment and translated the phases of all other time points accordingly so as to maintain their phase differences with respect to each other. This allowed us to compare the deviation of SVD inferred phases with respect to the true phases. As shown in Figure S9, the performance of our supervised algorithm was almost always better in comparison to the unsupervised approach using SVD.

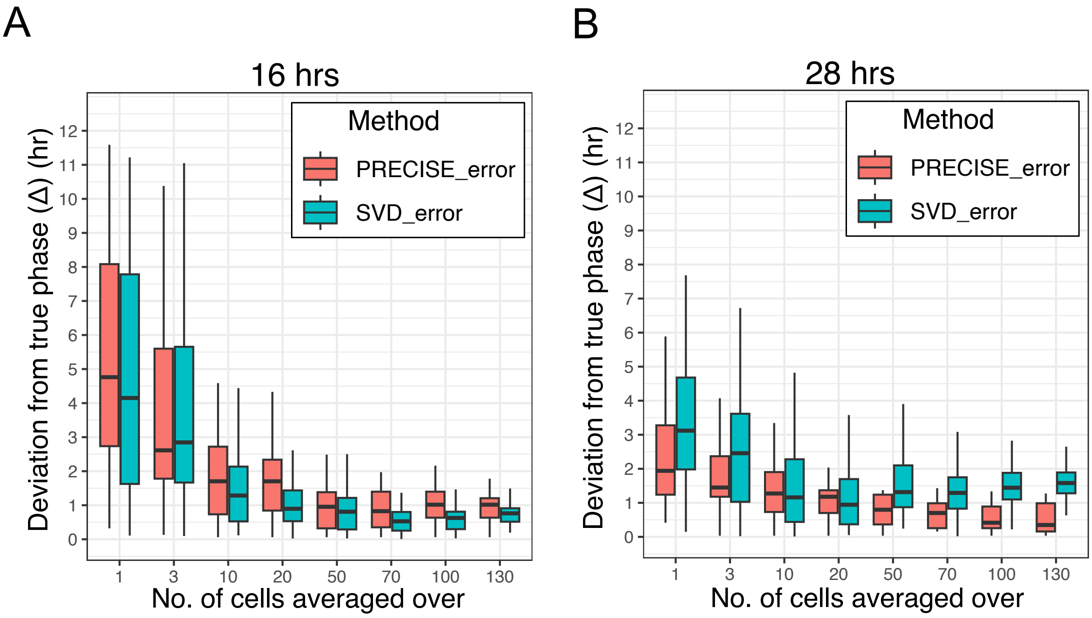

Figure S9: Comparison of deviations in phase inference using PRECISE and SVD for the different time points in the NIH3T3 test dataset.

#### 2. SVD based Circadian phase estimation with increasing gene numbers.

To explore whether the inability of clustering single cells using SVD can be a result of using only 4 genes, we performed a new experiment where we measured the expression of 6 genes namely – *Bmal1, Nr1d1, Nr1d2, Tef, Cry1* and *Nifl3* at 6 different time post Dexamethasone resetting as mentioned in Section S1. We first visualized the negative binomial distribution of the single-cell spot counts for each of the different genes over the different time point. As shown in Figure S10A each of the different genes measured demonstrated significant overlap at the different time points. For this dataset we performed SVD based circular phase inference when increasing number of clock genes were used – 3, 4, 5 and 6 clock genes. As described previously we created $3 \times 300$ matrix whose entries were expression values of 3 indicated clock genes (*Bmal1, Cry1* and *Tef*) in 50 randomly chosen single cells from each of the 6 different time points. We normalized the entries of the matrix as described in Section S6 such that the mean and variance for the expression of the clock genes over the different time points was 0 and 1 respectively. This matrix was then subjected to SVD and the projection of the samples onto the first 2 eigengene space was obtained. We repeated the procedure for increasing number of genes (4, 5 and 6 genes) and observed that at single-cell resolution the various clusters remained mixed as shown in Figure S10B. This suggests that using more number of genes for phase inference is unlikely to affect our results.

In order to further understand whether increasing the number of genes facilitates circadian phase inference at single-cell resolution, we estimated the coefficient of variation (COV) of the different genes we measured at each time point. The values are listed below in Table S5. When we calculated the mean COV over the different time post Dex-resetting for each gene we observed that *Nr1d2* is the least noisy among the genes that we measured. Hence for the next set of simulations we used the negative binomial distribution parameters (r - 'number of successes' and p - 'probability of success') obtained for *Nr1d2* gene.

| Gene | 08 hrs | 12 hrs | 16 hrs | 20 hrs | 24 hrs | 28 hrs |
|---|---|---|---|---|---|---|
| *Bmal1* | 0.78 | 0.64 | 0.82 | 0.95 | 1.22 | 0.9 |
| *Cry1* | 0.73 | 1.84 | 0.75 | 0.63 | 0.57 | 0.56 |
| *Tef* | 0.91 | 1.33 | 0.95 | 0.82 | 0.81 | 0.81 |
| *Nr1d1* | 2.8 | 1.06 | 1.04 | 0.91 | 1.55 | 1.07 |
| *Nr1d2* | 0.81 | 1.27 | 0.61 | 0.65 | 0.81 | 0.71 |
| *Nifl3* | 1.75 | 0.72 | 0.83 | 1.07 | 0.9 | 1.47 |

Table S5: Coefficient of Variation of single-cell spot distributions of 6 core-clock genes over different times post Dex-resetting.

We then computationally generated single-cell RNA counts for a range of different gene numbers (1 to 154) from negative binomial distributions, using parameter values of *Nr1d2* over the different time points, but with different phases (3 such simulated genes are shown in Figure S10A). We obtained the SVD projection by following the same methodology previously described for the single cells characterized by the RNA counts of the various measured and simulated genes onto the first 2 eigengene space. As shown in Figure S10C, for two representative cases (number of genes = 6 and 20), the single cells failed to cluster separately. In order to more systematically understand the number of genes required to get accurate clustering, we calculated the purity of the clusters as a function of the number of genes used to obtain the SVD projection (main text Figure 4). We performed k-means clustering based on the euclidean distances between the points in the eigengene space. To perform the k-means clustering we utilized the `Kmeans` function in the `amap` package in R with the arguments : centers = 6, method = "euclidean", nstart = 25, iter.max = 200. We next measured the purity of each cluster. For each cluster, we identified the most frequent time point among the cells it contained. We then calculated the proportion of cells in the cluster that belonged to this dominant time point — this value represents the cluster's purity. After computing the purity for all clusters, we took the average to obtain an overall measure of clustering quality. This metric will range between a value of 0 to 1, where 1 represents that each cluster has cells from a particular time point only and hence the cells from the different time points are well separated. We repeated this procedure in order to understand how the number of genes affects clustering quality. Since we are randomly generating these gene counts and also the phase differences existing between the simulated genes could affect the performance, we repeated the entire procedure 100 times and then plotted the mean trajectory obtained over the 100 runs as a function of number of genes in main text Figure 4G.

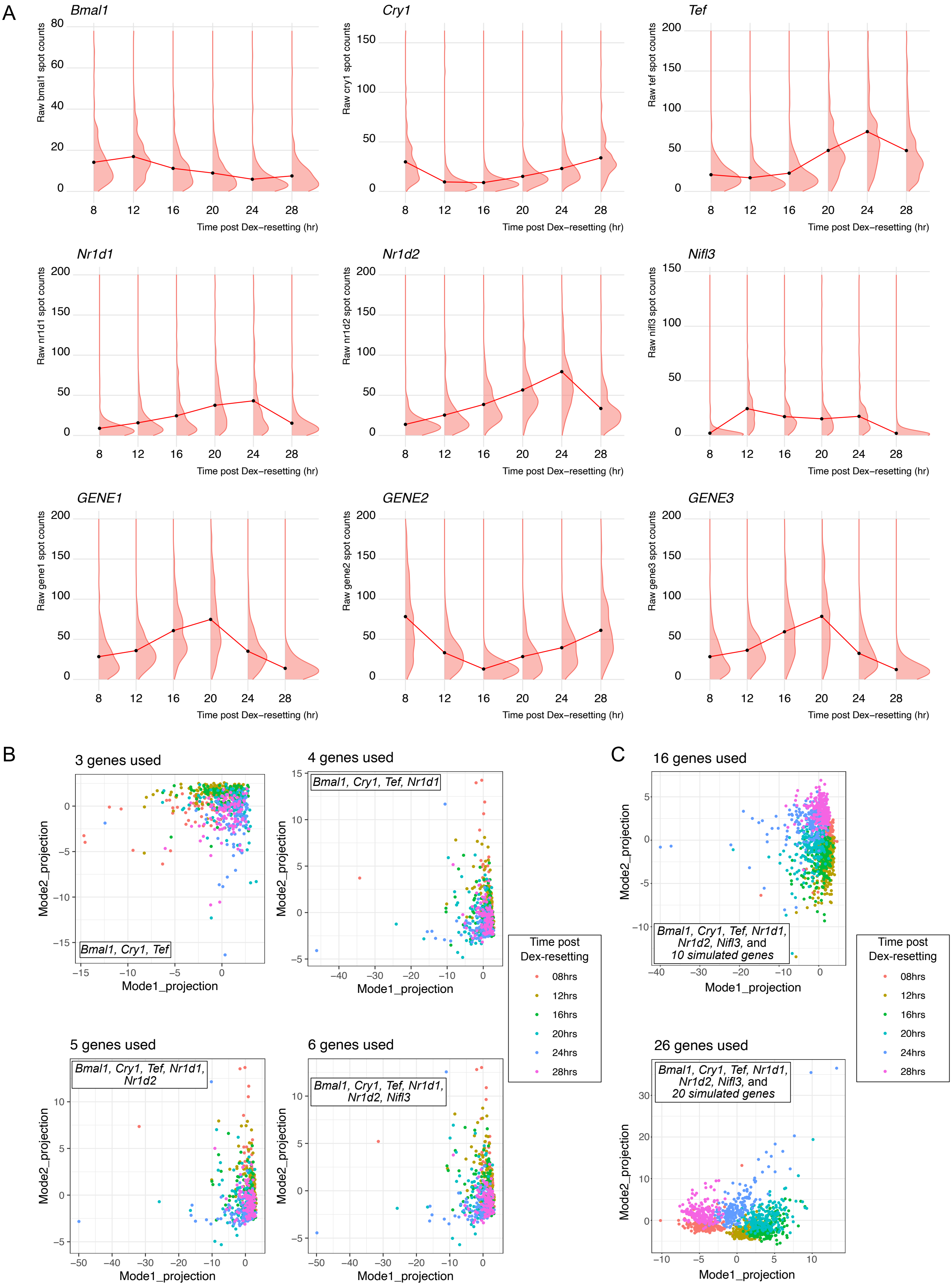

Figure S10: (A)Distributions of raw counts of different clock gene – *Bmal1, Nr1d1, Nr1d2, Tef, Cry1* and *Nifl3* along with the distributions for some of the simulated genes. Note the strongly overlapping distributions generated for the simulated genes.

Figure S10: (B) Increasing number of clock genes used to perform SVD-based phase inference. We observed that the cells from different clusters failed to separate and no elliptical structure was visible. (C) Simulating increasing number of genes with noise level as observed in experiments to understand to understand the role of gene number in single-cell phase inference. We observed that even with increasing number of genes (Top: 16 genes in total and Bottom: 26 genes in total) we did not observe a elliptical structure arise and also cells from different circadian phases clustered together.

We further checked if reduction of COV of the simulated genes (which would represent low noise core clock or clock-controlled genes) could lead to reduction in number of genes required for accurate clustering. In order to reduce the existing COV for each of the negative binomial distributions at different time points of the *Nr1d2* gene, we performed the following adjustment:

If $COV_i$ and $COV_f$ represent the initial and final COVs of the distributions respectively, and we desire the final COV to be 30% of the initial COV, then we have:

$$COV_f = 0.3 * COV_i \tag{20}$$

which implies that $\mu_f = (\mu_i/0.3)$, where $\mu_i$ and $\mu_f$ are the initial and final means of the distribution.

In the parameterization where the negative binomial distribution is characterized by number of successes (denoted by $r$) and the probability of success in each trial (denoted by $p$), the mean of the distribution is denoted by

$$\mu = \frac{r(1-p)}{p} \tag{21}$$

In this case, to update $\mu_i$ to $(\mu_i/0.3)$ we simply change the parameter $r$ to $(r/0.3)$.

Hence we updated the value of the parameter $r$ to $r/0.3$ at each of the different time points. While theoretically we expect a decrease of $\sim$ 70% in the COV, we sampled 300 random numbers using the updated parameters and observed an overall decrease of $\sim$50% in the COV. We simulated low-noise genes using the updated parameter values and repeated 100 runs of the simulation to obtain the mean cluster purity as a function of number of genes. As shown in main text Figure 4, in comparison to high-noise scenario (original COV), reducing the noise (by decreasing the COV) did not significantly affect our clustering results. We still required about 50 genes to obtained a cluster purity of 90%.

### S8 Neural network based circadian phase inference.

Recent work has demonstrated the superiority of autoencoder architecture of neural networks in inferring circadian phase from expression data of different genes [9, 10]. We implemented a similar autoencoder neural network to infer circadian phases for both single-cell gene expression dataset as well as when average expression is utilised as input.

As shown in Figure S11 A, we first used SVD to obtain a lower rank approximation to our full gene $\times$ cell data matrix comprising gene expression of all the clock genes *Bmal1*, *Nr1d1*, *Nr1d2* and *Tef*. After spectral decomposition of the full data matrix into its constitutent 3 matrices, we kept only the first two singular values and manually set the rest of the singular values to zero. This leads to de-noising of the data matrix thus causing cleaner oscillations to emerge. This reconstructed, reduced rank matrix is then used to train the autoencoder neural network. Autoencoders can be divided into parts – the encoder that learns a low dimensional projection of the input and the decoder that reconstructs the input from this low dimensional representation. Taking inspiration from recent studies, we implemented an autoencoder with a circular bottleneck [9] to infer circadian phases of samples. Our encoder consists of four layers – (1) the input layer consisting of 4 neurons (each neuron inputs the expression value of a single clock gene), (2) hidden layer consisting of 128 linear neurons that learn a high dimensional representation (entry of each of 128 neurons is a linear combination of the values of the 4 neurons in the previous layer), (3) hidden layer consisting of 2 linear neurons (entry of both the neurons is a linear combination of the values of the 128 neurons in the previous layer), and lastly (4) the circular bottleneck, where the values of the 2 neurons in the last layer are projected onto a circle and the angular position of the resulting point represents the circadian phase of the sample whose gene expression vector is the input. Our decoder consisting three layers with dimensions as mentioned in Figure S10, reconstructs the input $4\times$ 1 gene expression vector from the low dimensional representation.

Training the above mentioned autoencoder neural network entails learning the weights of the different linear layers included, and was achieved by reducing the mean squared error (MSELoss) between the input and the reconstructed output from the decoder. Once a trained autoencoder was obtained, the circadian phase values were determined using the trained encoder's circular bottleneck layer. We assembled this autoencoder using the `pytorch` package in Python [11].

Implementation of the SVD-autoencoder once again demonstrated that at the level of single cells phase inference was not possible (Figure S11 B). The Rayleigh statistic for cells from any single time point was low, suggesting a wide distribution on the circle. In contrast however, when expression was averaged over cellular groups, phase inference was perfectly feasible as can be seen from the Rayleigh statistics in Figure S11 C. Each of the samples showed Rayleigh statistics close to 1, suggesting clean clustering based on the true phases.

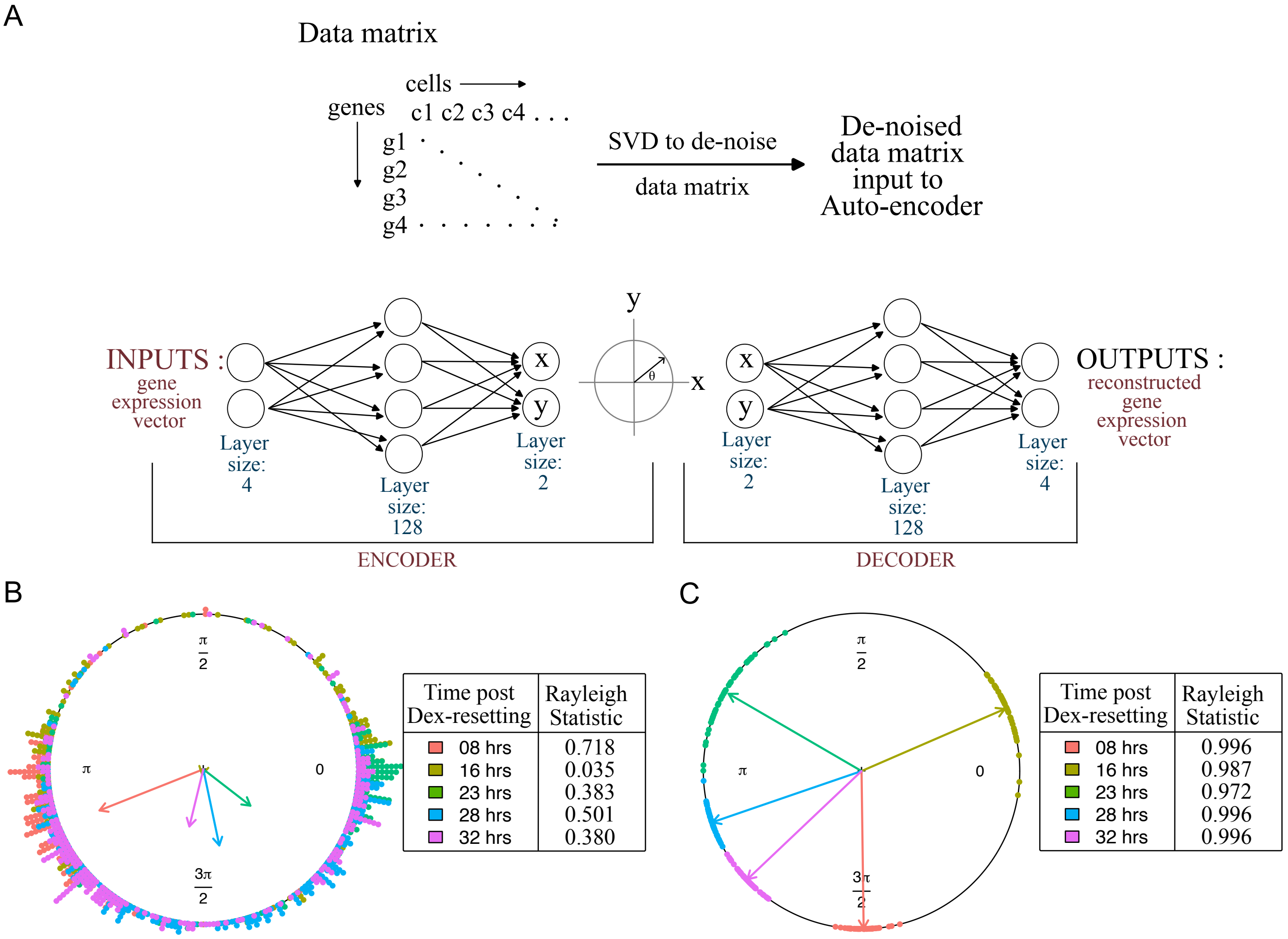

Figure S11: Autoencoder based inference of circadian phase is possible only after averaging. (A) Workflow for implementation of SVD followed autoencoder based circadian phase inference. (B) Phase inference performed at the level of single cells. The inferred phases are spread across the entire circle, suggesting transcriptional noise precludes phase inference at level of single cells. (C) Circadian phase inferred for samples comprising gene expression averaged over 100 cells. Coherent phase emerges suggesting the need for coarse-graining to define circadian phase in biological samples.

### S9 Circadian phase prediction across different cell lines.

To explore the possibility of across-cell lines predictions we trained PRECISE using NIH3T3 time course data as described in Section S6, and estimated the circadian phase from the absolute RNA counts in MLG cell lines (which were collected as described in Section S1). We followed a similar procedure as performed for NIH3T3 to explore the minimum number of cells and genes required for accurate across cell-line phase estimation.

As shown in Figures S12-13, expression values of three genes when averaged over a group of 50 randomly chosen cells allows for accurate phase inference in synchronized populations of MLG cells. As we did not obtain proper oscillations for *Bmal1* gene from MLG cell line dataset hence we did not use *Bmal1* gene to assign true phase and perform the phase estimation (Figure S13).

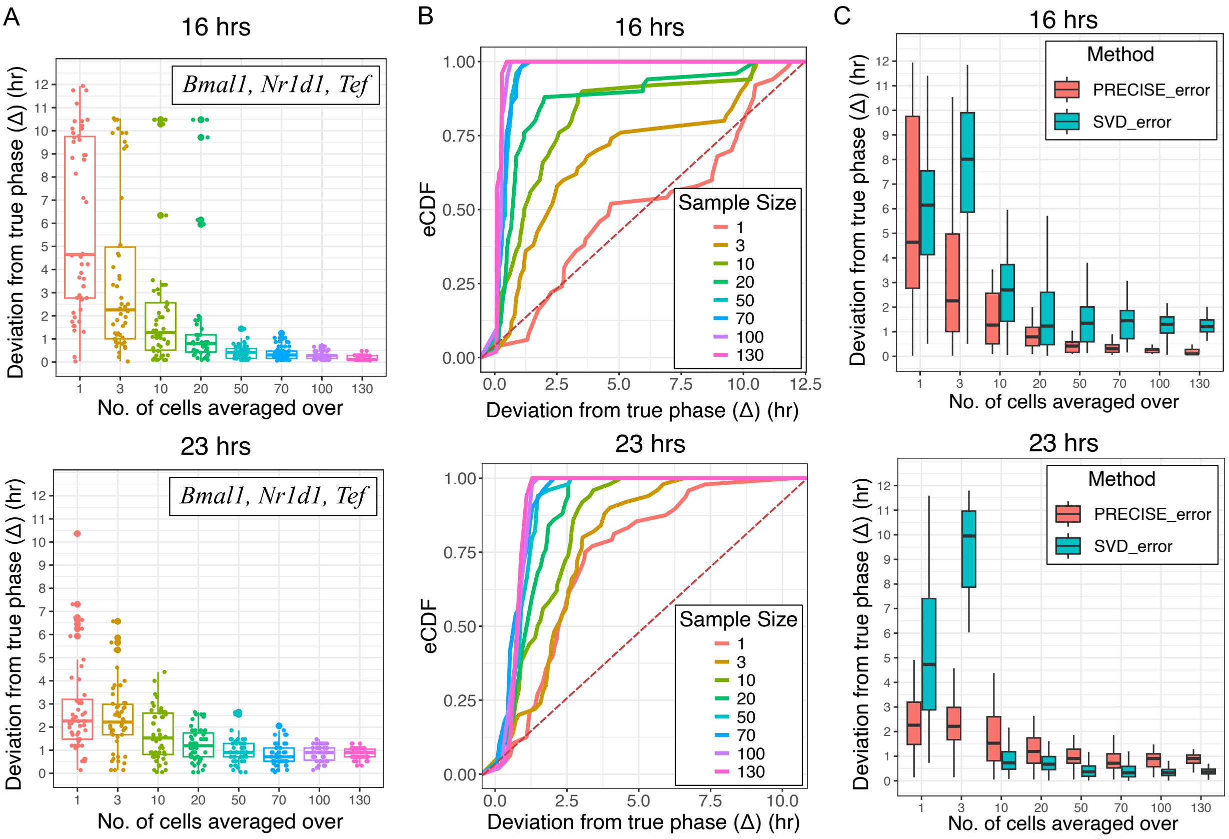

Figure S12: PRECISE based across cell-type phase prediction in Mouse Lung Fibroblasts (MLG). (A) Boxplot of deviations from true phase for the different time points. (B)eCDFs of deviations from true phase (C)Comparison of deviations in phase inference using PRECISE and SVD for the different time points in the MLG test dataset.

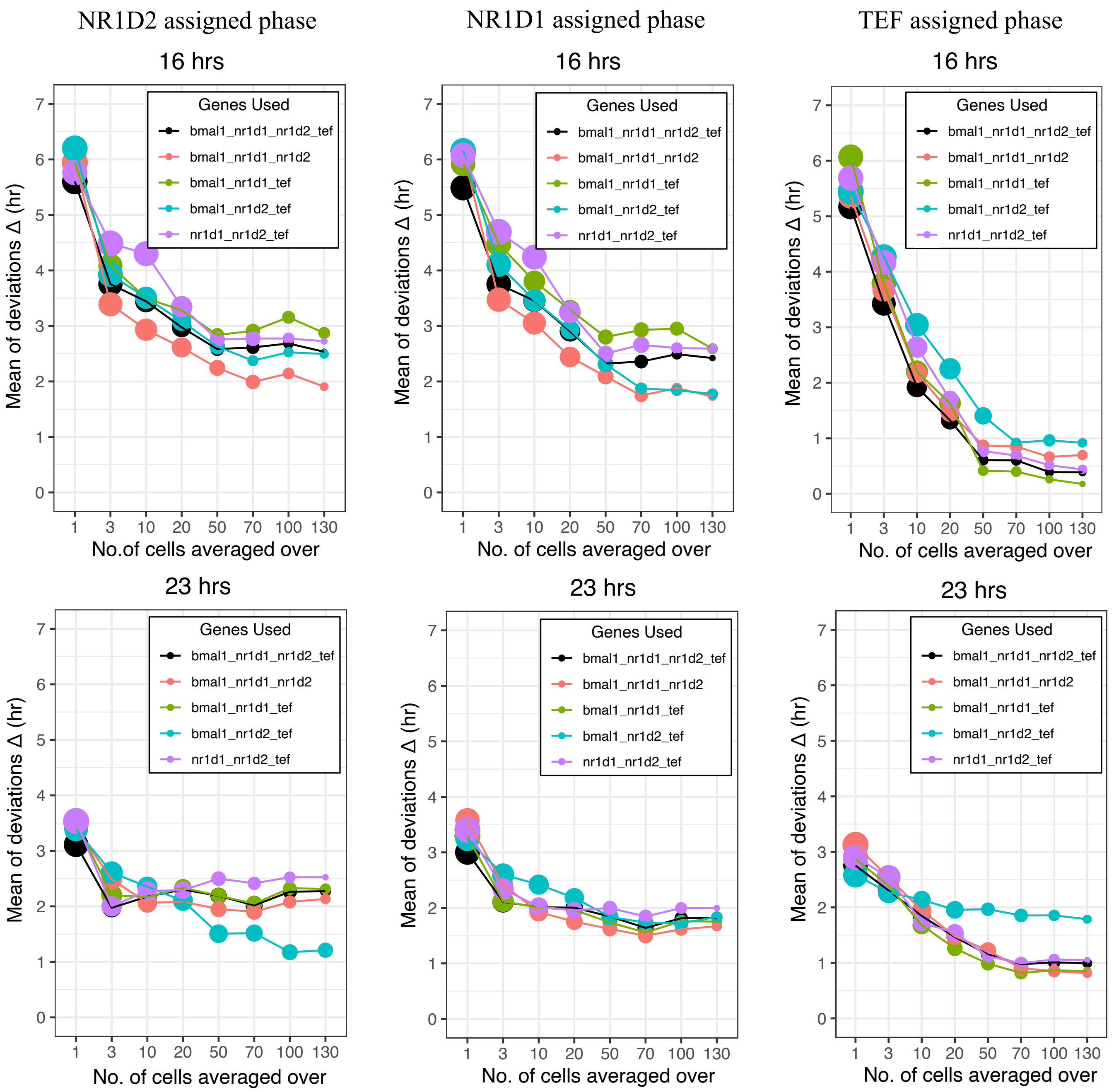

Figure S13: PRECISE-based across cell-type phase prediction in MLG cells when different genes were used to assign the true phases. For each of the different genes considered to assign true phase, we also compared the deviations across different possible combinations of three clock genes and also four gene combination across the three different test time points included in MLG data.

### S10 Cell-state discovery algorithms

#### 1. UMAP

Uniform Manifold Approximation and Projection (UMAP) is a dimensionality reduction technique that can be used to visualize a high-dimensional data in a lower-dimensional representation [12]. We implemented UMAP on our single-cell data comprising the gene expression for all 4 clock genes in all cells belonging to various different test time points. Before executing UMAP, we log-transformed the data where expression values $g$ of any gene was replaced by the $\log(g+1)$ value. We then obtained the UMAP of the log-transformed dataset using the `umap` function in the `umap` library of R by using the default parameter settings [13]. The n_components parameter controls the dimension of the space to which the user wants to embed the data to. The default value of this parameter is 2 and hence, we project down the data to a two-dimensional space. This function outputs the coordinates of the samples included in the dataset in the lower 2D space. Annotating the single cells by the known true circadian phase revealed that UMAP was unable to clusters cells from a particular phase together and the results remained unchanged for different learning rates used for estimation as shown in main text Figure 5 and Figure S14. We also repeated the UMAP based dimensionality reduction for a dataset comprising of 50 samples per time point where each sample is the average gene expression over 70 cells from that time point. Upon averaging the gene expression UMAP based dimensionality reduction also identified 5 distinct clusters as shown in main text Figure 5.

#### 2. t-SNE

t-SNE (t-distributed Stochastic Neighbor Embedding) is an unsupervised non-linear dimensionality reduction technique for data visualization [14]. To implement t-SNE on our log-transformed single-cell dataset described in the last section, we used the `Rtsne` function in the `Rtsne` library of R with the parameter values : check_duplicates = FALSE, initial_dims = 4 [15]. This function also projects the data onto a 2D space. Similar to UMAP, t-SNE failed to segregate the single cells based on their true circadian phase. However when t-SNE was implemented for 50 samples per time point where each sample is the average gene expression over 70 cells from the time point, clear segregation of the samples based on known true phase was observed (main text Figure 5).

#### 3. SEACells

Besides UMAP and t-SNE we also implemented a metacell identification algorithm that has been recently developed [16]. 'Metacells' represent distinct cell-states, where the variability within a metacell arises from technical rather than biological sources [17]. We implemented SEACells to identify 5 metacells using all 4 clock genes in our single-cell dataset. We log-transformed the data and obtained the metacells and assignment of the different single cells to their respective metacells. The value of the argument n_waypoint_eigs within the `SEACells.core.SEACells` function represents the number of eigenvectors (obtained by performing PCA on the input data matrix) to be considered while generating the metacells. While in the main text Figure 5 we included the metacell identification using 3 eigenvectors, we checked the results by varying the number of eigenvectors from 2 to 6. In each case as shown in Figure S14 we did not obtain pure metacells comprising of cells from a single time point. Similarly we also obtained metacells from our averaged dataset mentioned in the previous sections for the different number of eigenvectors considered.

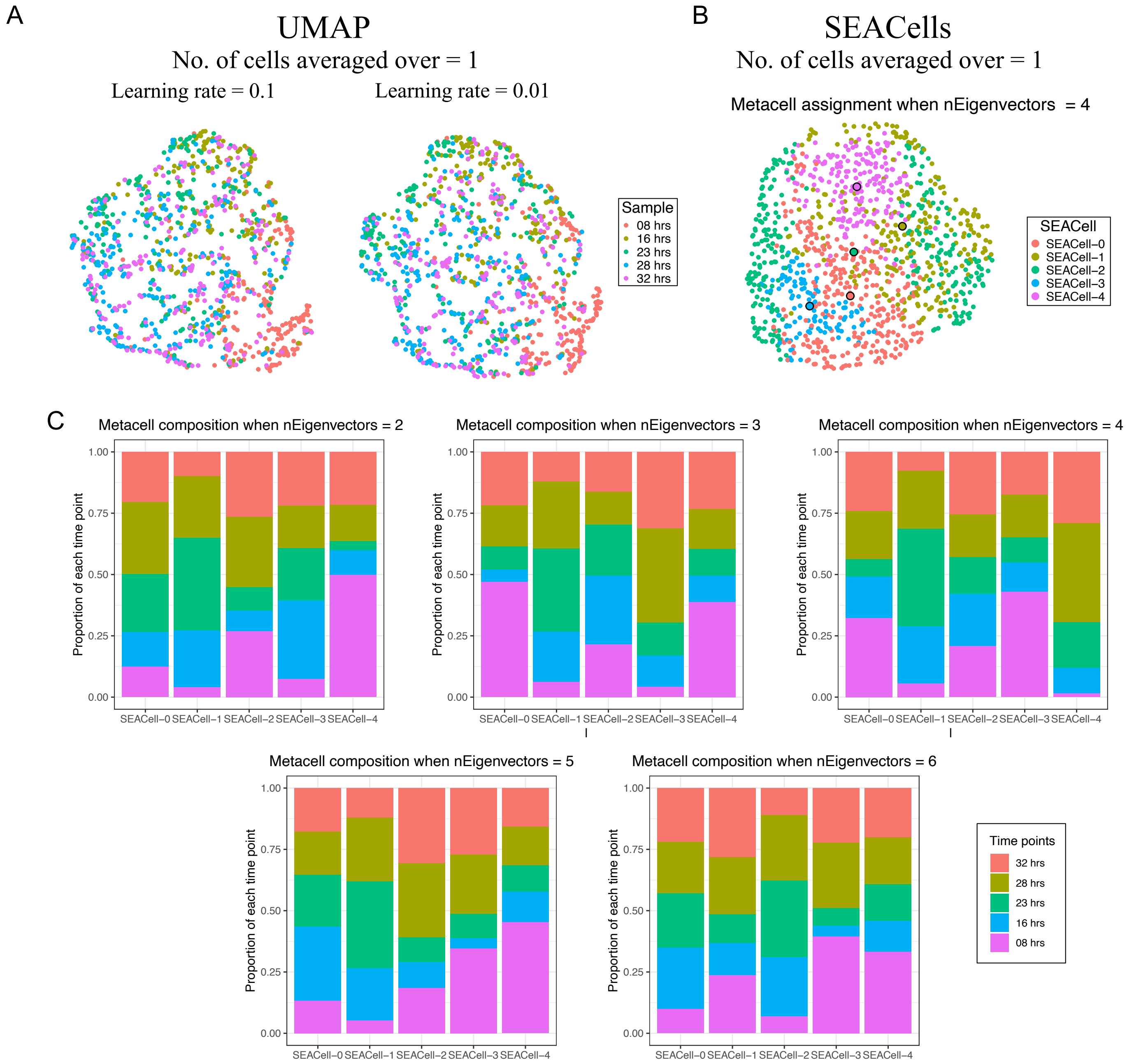

Figure S14: Different parameter runs for the cell-state discovery algorithms. (A) Lower learning rate in UMAP runs failed to cluster single cells into appropriate clusters based on their circadian phase. (B) The SEACells-based metacell assignment of single cells when 4 eigenvectors were considered. (C) Metacell composition for different SEACells runs using different numbers of eigenvectors.

In order to intuitively understand what enabled accurate circadian phase assignment for averages over cell groups as opposed to single cells, we visualized the distributions of raw spot counts for the clock genes across the various time points as shown in main text Figure 5E-F and S15. While the NB distributions of gene expression at the single-cell level were strongly overlapping, the approximately Gaussian distributions obtained upon averaging were well separated, thus allowing for accurate phase prediction.

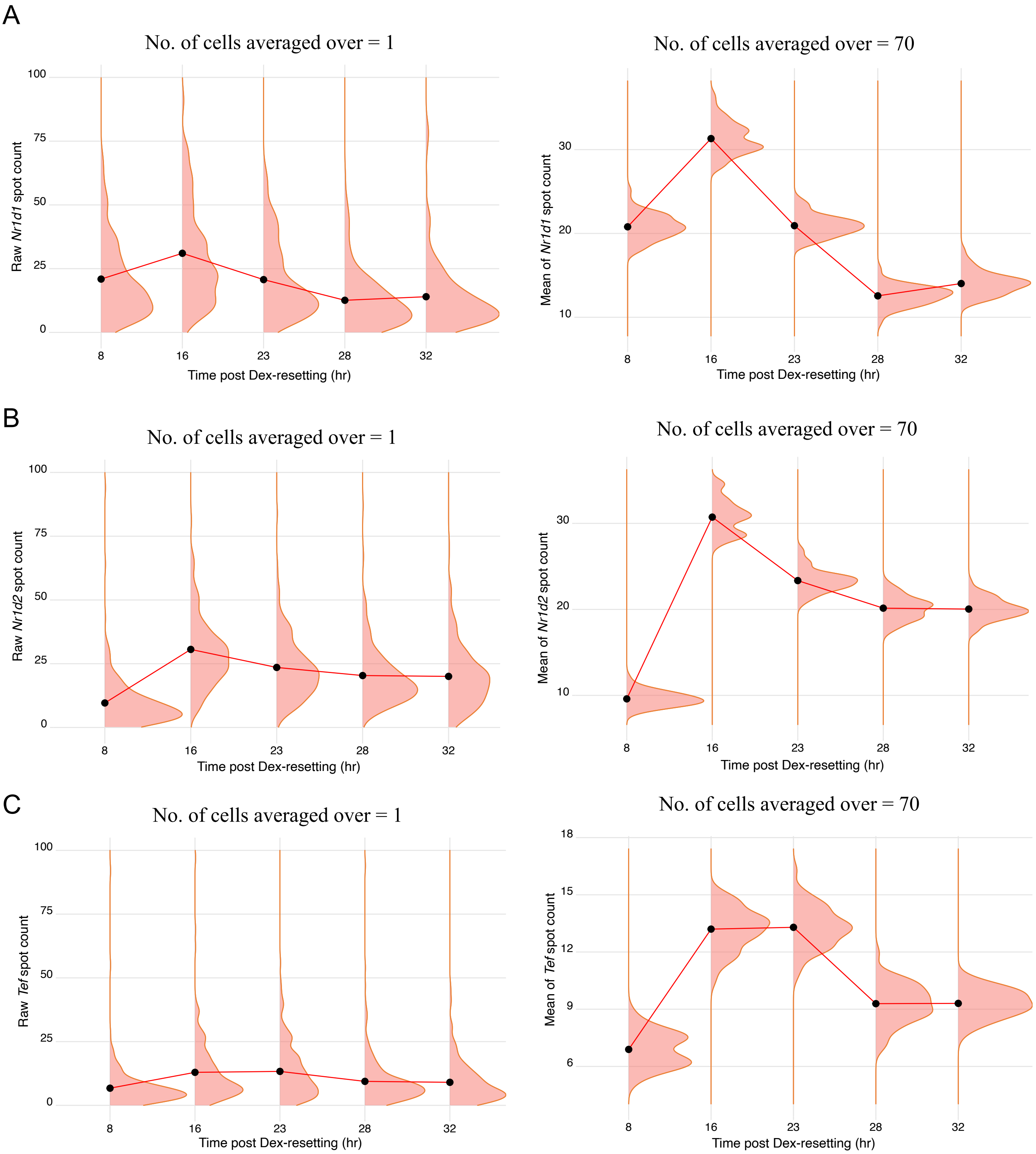

Figure S15: Distributions of raw and mean spot counts for the different clock genes in single cells across the 5 test time points post Dex-resetting. Distributions for *Bmal1* are shown in the main text Figure 5E-F.

### S11 Effect of sampling noise on circadian phase inference.

Commonly used multiplexed gene expression measurement techniques such as smFISH or scRNA-seq suffer from varying degrees of sampling noise, where every RNA molecule has a finite probability ($< 1$) of being captured and detected. Typically capture efficiencies tend to be much lower for sequencing based methods (such as scRNA-seq) as compared to imaging based methods (smFISH or MER-FISH). Here we asked whether and how much this added layer of variability contributes to our finding that circadian phases cannot be identified at single-cell resolution.

#### 1. Deriving the capture efficiency in SABER-FISH experiments from colocalization.

Colocalisation assays in smFISH aim to understand the specificity of binding of probes to the target molecule (say RNA). Two probe-sets, Probe 1 and Probe 2 are targeted to the same RNA and then the extent of colocalisation is measured.

In this scenario, let $p_1$ be the probability that any given RNA in the cell is bound by Probe 1. Similarly, define $p_2$ to be the probability that any RNA is bound by Probe 2. Then the following expressions denote the kinds of smFISH spots observed:

- $p_1$ = fraction of total RNA bound by probe 1
- $p_2$ = fraction of total RNA bound by probe 2
- $p_1 p_2$ = fraction of total RNA bound by both probe1 and probe2 (colocalised probes)
- $(1 - p_1)(1 - p_2)$ = fraction of total RNA not bound by any probe (hence not observed in the experiment).

Total sampled RNA (by either probe) will be given by:

$$\text{Sampled\_RNA} = p_1 + p_2 - p_1 p_2. \tag{22}$$

Hence the fraction of colocalized spots can be expressed as:

$$\text{Colocalisation} = \frac{p_1 p_2}{p_1 + p_2 - p_1 p_2}. \tag{23}$$

Poor sampling or capture probabilities would mean that $p_1 \to 0$ and $p_2 \to 0$. We calculated the change in colocalisation as a function of the values of $p_1$ and $p_2$ as shown below. With increase in capture efficiency of both probes, the observed colocalization increases (Figure S16 A). For reference, the lower limit of colocalization we observe in our experiments, $\sim 90\%$, is marked (Figure S16 B) – showing that to obtain such a high value of colocalization, both capture probabilities $p_1$ and $p_2$ have to be equal to $\sim 95\%$. This implies that for any probe-set, only 5% of the total RNA transcripts in a single-cell will be missed by our experimental protocol.

#### 2. Simulations to explore the effect of sampling noise on clustering and circadian phase identification.

We generated synthetic distributions of single-cell RNA counts for four different genes, assuming two distinct phases for simplicity. We considered the situation when the underlying distributions are overlapping in the two distinct phases due to biological noise. We utilized the NB parameters derived from fits to our training datasets and considered the time points 42 and 46 hours post Dex-resetting to represent the two states or circadian phases. For each of the two phases, we sampled 200 individual cells each characterised by a vector of absolute counts of 4 genes sampled from the underlying distribution. These counts per cell that we generated were considered the true counts or the actual RNA numbers in the different cells. From these true counts, we sampled with sampling probabilities less than 1 as indicated in the figures and performed UMAP on the resulting data. After generating the sampled data, we also performed averaging over small groups of 30 randomly chosen cells and repeated the procedure 50 times to generate 50 averaged samples per phase.

From Figure S16, we see that regardless of low or high sampling probability, if the underlying distributions are overlapping then we do not obtain well separated clusters in the low dimensional projection when using single-cell gene expression as input. However upon averaging, we recovered the true circadian phases using UMAP.

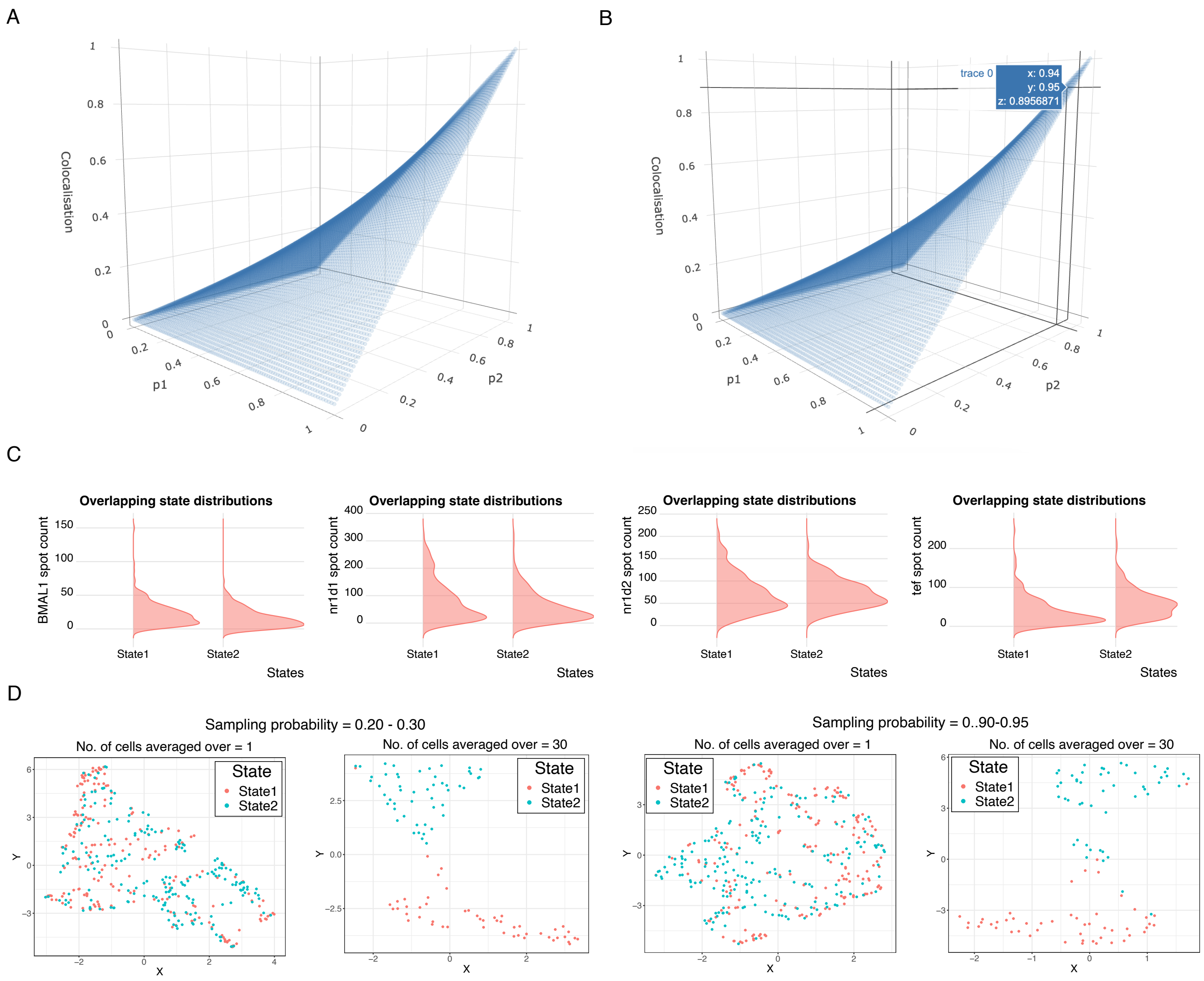

Figure S16: Effect of sampling noise on circadian phase inference. (A) Change in colocalisation as a function of individual sampling (capture) probabilities of two probe-sets in the context of a colocalisation smFISH experiment. (B) To reach $\sim 90\%$ colocalization, each probe-set must bind with $\sim 95\%$ capture probability. (C) Distribution of absolute RNA numbers when the underlying single-cell gene expression distributions of two distict circadian phases are overlapping. (D) (Left) UMAP projection of single cells and 50 averaged samples (single sample comprises expression averaged over 30 randomly chosen cells) when the sampling probability is low (0.20 - 0.30). (Right) UMAP projection of single cells and 50 averaged samples (single sample comprises expression averaged over 30 randomly chosen cells) when the sampling probability is low (0.90 - 0.95). The samples in D are coloured by true phase.

### S12 Spatial Phase Inference from a population of asynchronized NIH3T3 cells.

#### 1. Rayleigh statistic to measure phase coherence.

In order to quantify the spread in inferred phase from a single time point/spatial group, we calculated the Rayleigh's statistic. The Rayleigh's test for uniformity measures the spread in distribution of a circular variable. If the points are uniformly distributed around the circle then the value the statistic is close to 0, and the length of the mean resultant vector is small. However when the points are restricted to a specific region on the circle, the value of the statistic is close to 1 and the mean resultant length is also high. To calculate the length of the resultant vector and the Rayleigh's statistic we utilised the `rayleigh.test` function with the default parameters included in the `circular` package in R [8].

#### 2. Stitched images to acquire spatial data from an asynchronized population of NIH3T3 cells.

We seeded cells in a single four-chambered slide and allowed them to grow in the wells for sufficient time so as to remove spurious synchronisation that may occur during seeding. Following the SABER-FISH protocol as mentioned in Section S1, we obtained the counts of the four clock genes for the asynchronized population.

From our analysis on synchronised populations of cells we demonstrated that averaging gene expression over 20-50 cells was essential to obtain an accurate phase estimate. As mentioned in Section S1, imaging the cells at 60X magnification usually results in less than 20 cells per field of view. Hence to obtain sufficient number of spatially proximal cells to carry out the averaging process, we imaged cells using the stitched image setting in Nikon's NIS elements software. We imaged $2\times2$ grid images that were later stitched with 5% overlap. This procedure allows us to capture $\sim$ 45-60 cells per $2\times2$ grid. In this way we imaged 7 different spatial groups as mentioned in the main text.

#### 3. Calculating the distance between different spatial groups of cells.

Sufficiently spatially separated groups of cells are expected to represent distinct circadian phases. Hence we quantified the distances between the different spatial groups of cells we imaged, to ensure that within-group distances were significantly less than across-group distances. As shown in Figure S15 and described in the previous section, the cells belonging to a single spatial group were imaged as a 2×2 grid image. We obtained the centre coordinates of the image for each of the grid images (depicted by the blue circles in Figure S17) as well as the coordinates of the centre for the stitched image (shown by the red circle in Figure S17) with respect to the stage coordinate axes. To measure the intra-group distances, we calculated the euclidean distances between the centre coordinates of all possible combination of grid images (blue arrowhead-lines) within a single spatial group. Similarly inter-group distances were calculated as the euclidean distances between the centre coordinates of the stitched images across all different combinations of spatial groups (red arrowhead-lines). As shown in main text Figure 6, following the above method we observed that the inter-group distances were significantly higher than the intra-group distances, suggesting that our different spatial groups of cells were well separated compared to the cellular distances within each spatial group.

Post image analysis (as described in Section S2) of the asynchronized cellular population, we proceeded to infer PRECISE-based circadian phases for each of the different spatial groups separately when gene expression was averaged over groups of 20 cells present within a spatially proximal group of cells. As shown in main text Figure 6D at the population level the inferred phases were spread across the circle corroborating the fact that our population was truly asynchronized. However as shown in Figure 6D and S18, distinct coherent phases were observed for within each spatial group, supporting our expectation that even in an otherwise asynchronized population, the spatially proximal cells tend to be in similar phases.

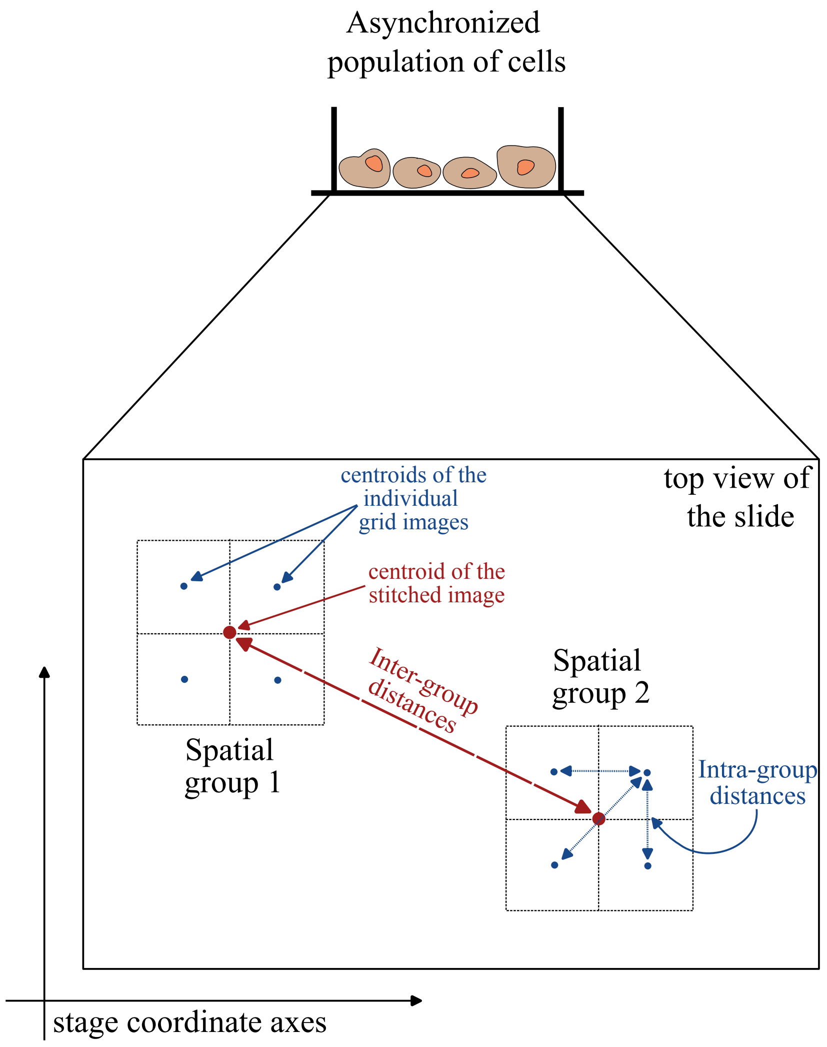

Figure S17: Overview of method to calculate the distances within and across different spatial groups of cells. Cells imaged within 2×2 grids represent spatially proximal cells.

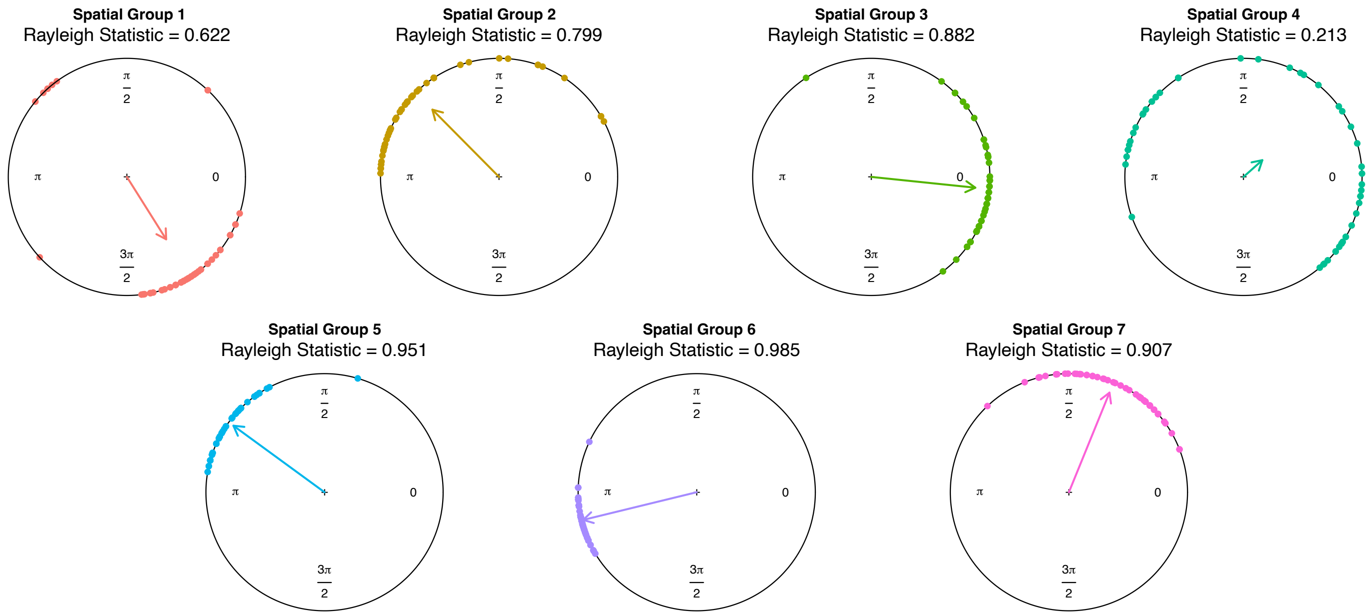

Figure S18: High phase coherence is observed within the different spatial groups as quantified by the high individual Rayleigh statistics. The only exception was spatial group 4.
